# Assessment of network module identification across complex diseases

**DOI:** 10.1101/265553

**Authors:** Sarvenaz Choobdar, Mehmet E. Ahsen, Jake Crawford, Mattia Tomasoni, Tao Fang, David Lamparter, Junyuan Lin, Benjamin Hescott, Xiaozhe Hu, Johnathan Mercer, Ted Natoli, Rajiv Narayan, The DREAM Module Identification Challenge Consortium, Aravind Subramanian, Jitao D. Zhang, Gustavo Stolovitzky, Zoltán Kutalik, Kasper Lage, Donna K. Slonim, Julio Saez-Rodriguez, Lenore J. Cowen, Sven Bergmann, Daniel Marbach

## Abstract

Identification of modules in molecular networks is at the core of many current analysis methods in biomedical research. However, how well different approaches identify disease-relevant modules in different types of gene and protein networks remains poorly understood. We launched the “Disease Module Identification DREAM Challenge”, an open competition to comprehensively assess module identification methods across diverse protein-protein interaction, signaling, gene co-expression, homology, and cancer-gene networks. Predicted network modules were tested for association with complex traits and diseases using a unique collection of 180 genome-wide association studies (GWAS). Our critical assessment of 75 contributed module identification methods reveals novel top-performing algorithms, which recover complementary trait-associated modules. We find that most of these modules correspond to core disease-relevant pathways, which often comprise therapeutic targets and correctly prioritize candidate disease genes. This community challenge establishes benchmarks, tools and guidelines for molecular network analysis to study human disease biology (https://synapse.org/modulechallenge).

## Introduction

Understanding the mechanisms and pathways underlying complex human diseases remains a difficult problem, hindering the development of targeted therapeutics. Complex diseases involve many genes and molecules that interact within context-specific cellular networks (Califano et al., 2012). These densely interconnected networks sense and propagate perturbations from genetic variants and environmental factors, giving rise to disease states that may be difficult to understand at the level of individual genes (Schadt, 2009). Indeed, it has become apparent that the majority of genetic variants underlying complex traits and diseases lie in noncoding regions of the genome where they presumably disrupt gene regulatory networks (Bonder et al., 2017; Westra et al., 2013), lending further support to the long-recognized importance of molecular network analysis for understanding disease biology (Ideker and Sharan, 2008; Vidal et al., 2011).

Experimental and computational techniques for mapping molecular networks, including physical interaction networks (e.g., protein-protein interaction, signaling and regulatory networks) as well as functional gene networks (e.g., co-expression and genetic interaction networks), have been a major focus of systems biology. Recent studies have further introduced comprehensive collections of tissue-specific networks (Greene et al., 2015; Marbach et al., 2016). Network-based approaches are now widely used for systems-level analyses in diverse fields ranging from oncology (Tsherniak et al., 2017) to cell differentiation (Cahan et al., 2014; Ciofani et al., 2012). A key problem in biological network analysis is the identification of functional units, called modules or pathways. It is well known that molecular networks have a high degree of modularity (i.e., subsets of nodes are more densely connected than expected by chance), and that the corresponding modules often comprise genes or proteins that are involved in the same biological functions (Hartwell et al., 1999). Moreover, biological networks are typically too large to be examined and visualized as a whole. Consequently, module identification is often a crucial step to gain biological insights from network data (Chen et al., 2008; Huttenhower et al., 2009; Pe’er et al., 2001).

Module identification, also called community detection or graph clustering, is a key problem in network science for which a wide range of methods have been proposed (Fortunato and Hric, 2016). These methods are typically assessed on *in silico* generated benchmark graphs (Girvan and Newman, 2002). However, how well different approaches uncover biologically relevant modules in real molecular networks remains poorly understood. Crowdsourced open-data competitions (known as challenges) have proven an effective means to rigorously assess methods and, in the process, foster collaborative communities and open innovation. The Dialogue on Reverse Engineering and Assessment (DREAM) is a community-driven initiative promoting open-data challenges in systems biology and translational medicine (http://dreamchallenges.org). DREAM challenges have established standardized resources and robust methodologies for diverse problems, including the inference of gene regulatory and signaling networks (Hill et al., 2016; Marbach et al., 2012). But, so far there has been no community effort addressing the downstream analysis of molecular networks.

Here we present the results of the Disease Module Identification DREAM Challenge (Fig. 1). The aim of this challenge is to comprehensively assess module identification methods across diverse molecular networks. Six research groups contributed unpublished gene and protein networks and over 400 participants from all over the world developed and applied module identification methods. In order to test the hypothesis that data integration across multiple types of networks enables more refined module predictions, teams were challenged to identify disease-relevant modules both within individual networks (Sub-challenge 1) and across multiple layered networks (Sub-challenge 2). In the final round, 75 submissions, including method descriptions and code, were made across the two sub-challenges, providing a broad sampling of state-of-the-art methods. We employed a novel approach to assess the performance of these methods based on the number of discovered modules associated with complex traits or diseases. In this paper, we discuss the top-performing approaches, show that they recover complementary modules, and introduce a method to generate robust consensus modules. Finally, we explore the biology and therapeutic relevance of trait-associated network modules.

**Figure 1:**
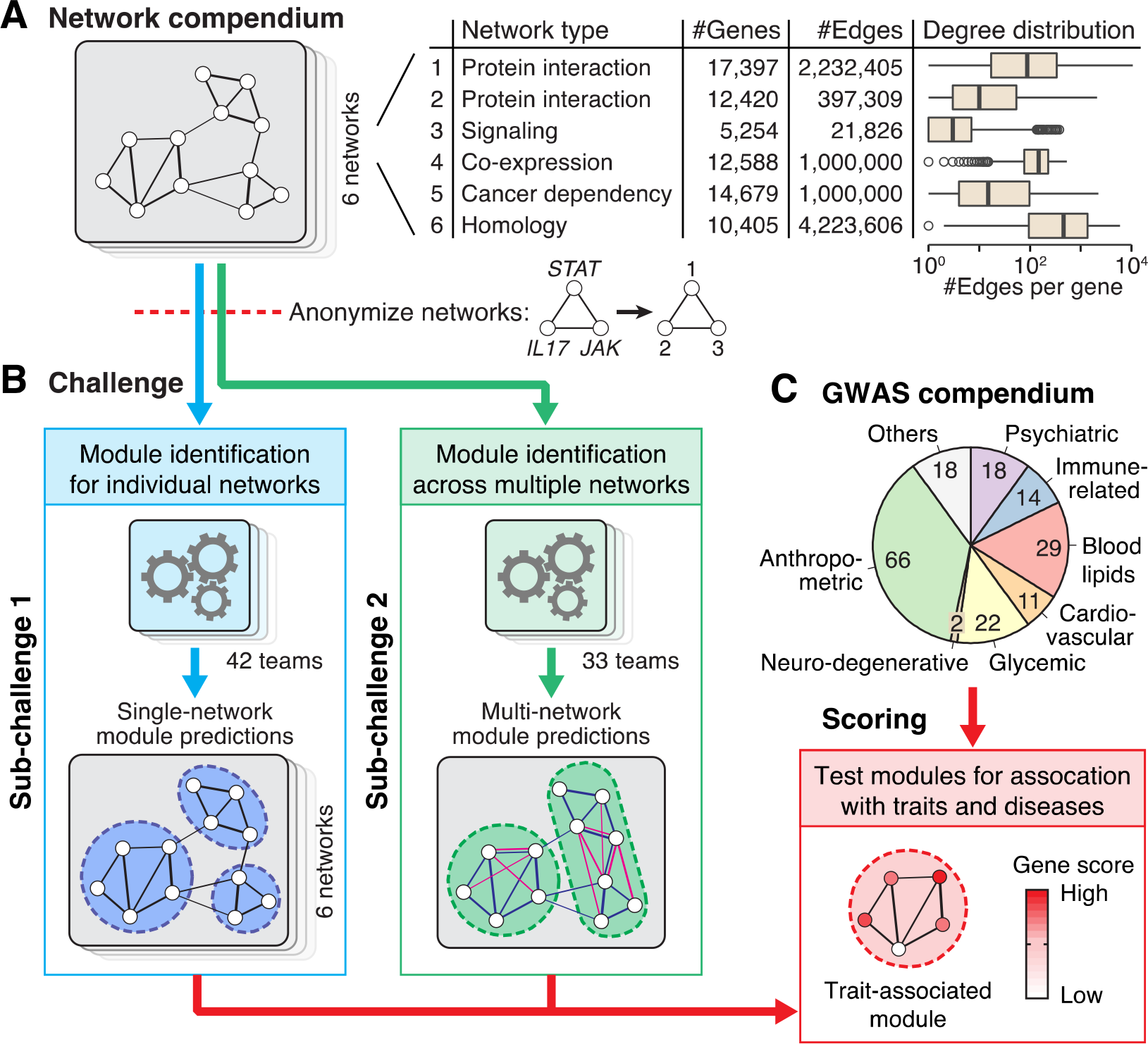
The Disease Module Identification DREAM Challenge. We launched an open-participation community challenge, where teams competed to predict groups of functionally related genes (i.e., modules) within diverse molecular networks. **(A)** The challenge comprised six networks, including protein-protein interaction, signaling, co-expression, cancer dependency, and homology-based gene networks. As the networks were all unpublished, we could anonymize them by removing the gene labels. This prevented participants from using existing knowledge of gene functions, thus enabling rigorous, blinded assessment. (Throughout the paper, boxplot center lines show the median; box limits, upper and lower quartiles; whiskers, 1.5x interquartile range; points, outliers.) **(B)** The aim of the challenge was to identify disease-relevant modules within the provided networks. Teams could participate in either or both sub-challenges: 42 teams predicted modules for individual networks (Sub-challenge 1) and 33 teams predicted integrated modules across multiple networks (Sub-challenge 2). **(C)** The submitted modules were tested for association with complex traits and diseases using a comprehensive collection of 180 GWAS datasets. The final score for each method was the number of trait-associated modules that it discovered. Since GWAS are based on data completely different from those used to construct the networks, they can provide independent support for biologically relevant modules.

All challenge data, including the networks, GWAS datasets, team submissions and code are available as a community resource at https://www.synapse.org/modulechallenge.

## Results

### A crowdsourced challenge for empirical assessment of module identification methods

We developed a panel of diverse, human molecular networks for the challenge, including custom versions of two protein-protein interaction and a signaling network extracted from the STRING (Szklarczyk et al., 2015), InWeb (Li et al., 2017) and OmniPath (Türei et al., 2016) databases, a co-expression network inferred from 19,019 tissue samples from the GEO repository (Subramanian et al., 2017), a network of genetic dependencies derived from genome-scale loss-of-function screens in 216 cancer cell lines (Cowley et al., 2014; Tsherniak et al., 2017), and a homology-based network built from phylogenetic patterns across 138 eukaryotic species (Li et al., 2018, 2014) (Methods). We included networks derived from different types of protein and gene associations, which also vary in their size and structural properties, to provide a heterogeneous benchmark resource and to evaluate the relative strengths of different network types for identifying disease-relevant modules (Fig. 1A).

Each network was generated specifically for the challenge and released in anonymized form (i.e., we did not disclose the gene names and the identity of the networks). Using unpublished networks made it impossible for participants to infer the gene identities, thus enabling rigorous “blinded” assessment. That is, participants could only use unsupervised clustering algorithms, which rely exclusively on the network structure and do not depend on additional biological information such as known disease genes. It is important to benchmark unsupervised methods in this blinded setting, because they are often relied on in regions of the network for which a paucity of biological information is currently available.

We solicited participation in two types of module identification challenges (Fig. 1B). In Sub-challenge 1, solvers were asked to run module identification on each of the provided networks individually (single-network module identification). Thus, they were asked to submit one set of modules for each of the six networks. This is a typical problem in biomedical research, where one is often presented with a single network derived from a given dataset. In Sub-challenge 2, the networks were re-anonymized in a way that the same gene identifier represented the same gene across all six networks. Solvers were then asked to identify a single set of non-overlapping modules by sharing information across the six networks (multi-network module identification), which allowed us to assess the potential improvement in performance offered by emerging multi-network methods compared to single-network methods. In both sub-challenges, predicted modules had to be non-overlapping and comprise between 3 and 100 genes (modules with over one hundred genes are typically less useful to gain specific biological insights).

We developed a framework to empirically assess module identification methods based on the number of predicted modules that show significant association with complex traits and diseases (called trait-associated modules, Fig. 1C). To this end, predicted modules were scored on GWAS data using the Pascal tool (Lamparter et al., 2016), which takes into account confounders such as linkage disequilibrium within and between genes (Methods). Since we are employing a large collection of 180 GWAS datasets ranging over diverse disease-related human phenotypes (**Table S1**), this approach covers a broad spectrum of molecular processes.

In contrast to evaluation of module enrichment using existing gene and pathway annotations, where it is sometimes difficult to ascertain that annotations were not derived from similar data types as the networks, the GWAS-based approach provides an orthogonal means to assess disease-relevant modules.

The challenge was run using the open-science Synapse platform (Derry et al., 2012). Over a two-month period, teams could make repeated submissions and see their performance on a real-time leaderboard to iteratively improve their methods. The total number of leaderboard submissions per team was limited to 25 and 41 for the two sub-challenges, respectively. In the final round, teams could make a single submission for each sub-challenge, which had to include detailed method descriptions and code for reproducibility. The scoring of the final submissions was based on a separate set of GWAS data sets that were not used during the leaderboard round (Methods).

### Community-based collection of module identification methods

The community contributed 42 single-network and 33 multi-network module identification methods in the final round of the two sub-challenges. Single-network module identification methods are listed in Table 1, top-performing approaches are detailed in Methods, and full descriptions and code of all methods are available on the Synapse platform (https://www.synapse.org/modulechallenge). In the following sections we first discuss the single-network methods (Sub-challenge 1).

We grouped methods into seven broad categories: (i) kernel clustering, (ii) modularity optimization, (iii) random-walk based, (iv) local methods, (v) ensemble methods, (vi) hybrid methods and (vii) other methods (Fig. 2A, Table 1). While many teams adapted existing algorithms for community detection, other teams -- including the best performers -- developed novel approaches.

**Table 1.**
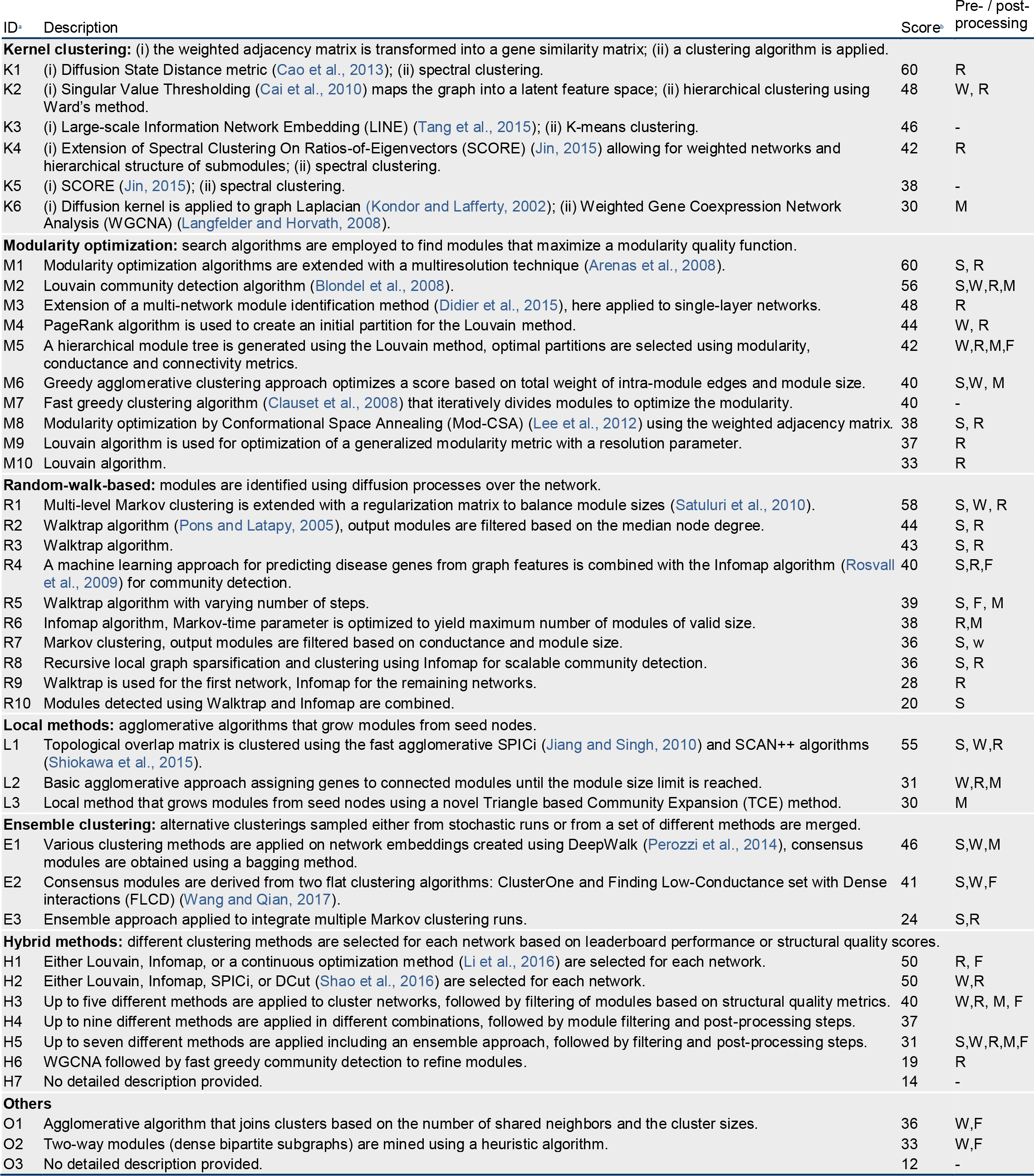
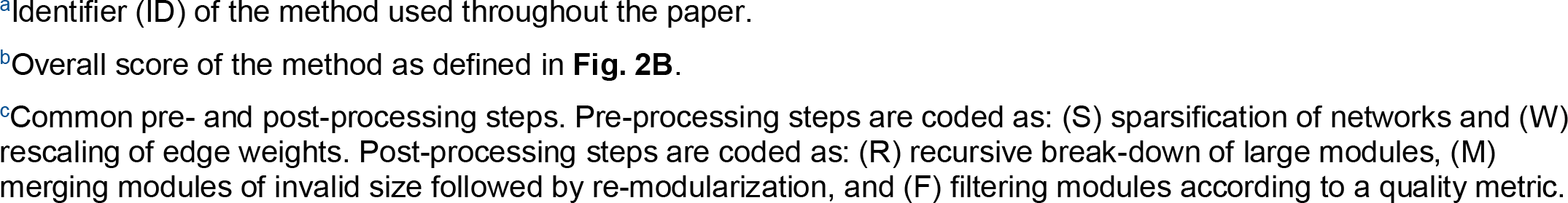
Module identification methods. The 42 module identification methods applied in Sub-challenge 1 grouped by category (see Fig. 2A).

| ID | Description | Score | Pre- / post-processing |
| --- | --- | --- | --- |
| **Kernel clustering:** (i) the weighted adjacency matrix is transformed into a gene similarity matrix; (ii) a clustering algorithm is applied. |  |  |  |
| K1 | (i) Diffusion State Distance metric (Cao et al., 2013); (ii) spectral clustering. | 60 | R |
| K2 | (i) Singular Value Thresholding (Cai et al., 2010) maps the graph into a latent feature space; (ii) hierarchical clustering using Ward's method. | 48 | W, R |
| K3 | (i) Large-scale Information Network Embedding (LINE) (Tang et al., 2015); (ii) K-means clustering. | 46 | - |
| K4 | (i) Extension of Spectral Clustering On Ratios-of-Eigenvectors (SCORE) (Jin, 2015) allowing for weighted networks and hierarchical structure of submodules; (ii) spectral clustering. | 42 | R |
| K5 | (i) SCORE (Jin, 2015); (ii) spectral clustering. | 38 | - |
| K6 | (i) Diffusion kernel is applied to graph Laplacian (Kondor and Lafferty, 2002); (ii) Weighted Gene Coexpression Network Analysis (WGCNA) (Langfelder and Horvath, 2008). | 30 | M |
| **Modularity optimization:** search algorithms are employed to find modules that maximize a modularity quality function. |  |  |  |
| M1 | Modularity optimization algorithms are extended with a multiresolution technique (Arenas et al., 2008). | 60 | S, R |
| M2 | Louvain community detection algorithm (Blondel et al., 2008). | 56 | S,W,R,M |
| M3 | Extension of a multi-network module identification method (Didier et al., 2015), here applied to single-layer networks. | 48 | R |
| M4 | PageRank algorithm is used to create an initial partition for the Louvain method. | 44 | W, R |
| M5 | A hierarchical module tree is generated using the Louvain method, optimal partitions are selected using modularity, conductance and connectivity metrics. | 42 | W,R,M,F |
| M6 | Greedy agglomerative clustering approach optimizes a score based on total weight of intra-module edges and module size. | 40 | S,W, M |
| M7 | Fast greedy clustering algorithm (Clauset et al., 2008) that iteratively divides modules to optimize the modularity. | 40 | - |
| M8 | Modularity optimization by Conformational Space Annealing (Mod-CSA) (Lee et al., 2012) using the weighted adjacency matrix. | 38 | S, R |
| M9 | Louvain algorithm is used for optimization of a generalized modularity metric with a resolution parameter. | 37 | R |
| M10 | Louvain algorithm. | 33 | R |
| **Random-walk-based:** modules are identified using diffusion processes over the network. |  |  |  |
| R1 | Multi-level Markov clustering is extended with a regularization matrix to balance module sizes (Satuluri et al., 2010). | 58 | S, W, R |
| R2 | Walktrap algorithm (Pons and Latapy, 2005), output modules are filtered based on the median node degree. | 44 | S, R |
| R3 | Walktrap algorithm. | 43 | S, R |
| R4 | A machine learning approach for predicting disease genes from graph features is combined with the Infomap algorithm (Rosvall et al., 2009) for community detection. | 40 | S,R,F |
| R5 | Walktrap algorithm with varying number of steps. | 39 | S, F, M |
| R6 | Infomap algorithm, Markov-time parameter is optimized to yield maximum number of modules of valid size. | 38 | R,M |
| R7 | Markov clustering, output modules are filtered based on conductance and module size. | 36 | S, w |
| R8 | Recursive local graph sparsification and clustering using Infomap for scalable community detection. | 36 | S, R |
| R9 | Walktrap is used for the first network, Infomap for the remaining networks. | 28 | R |
| R10 | Modules detected using Walktrap and Infomap are combined. | 20 | S |
| **Local methods:** agglomerative algorithms that grow modules from seed nodes. |  |  |  |
| L1 | Topological overlap matrix is clustered using the fast agglomerative SPICi (Jiang and Singh, 2010) and SCAN++ algorithms (Shiokawa et al., 2015). | 55 | S, W,R |
| L2 | Basic agglomerative approach assigning genes to connected modules until the module size limit is reached. | 31 | W,R,M |
| L3 | Local method that grows modules from seed nodes using a novel Triangle based Community Expansion (TCE) method. | 30 | M |
| **Ensemble clustering:** alternative clusterings sampled either from stochastic runs or from a set of different methods are merged. |  |  |  |
| E1 | Various clustering methods are applied on network embeddings created using DeepWalk (Perozzi et al., 2014), consensus modules are obtained using a bagging method. | 46 | S,W,M |
| E2 | Consensus modules are derived from two flat clustering algorithms: ClusterOne and Finding Low-Conductance set with Dense interactions (FLCD) (Wang and Qian, 2017). | 41 | S,W,F |
| E3 | Ensemble approach applied to integrate multiple Markov clustering runs. | 24 | S,R |
| **Hybrid methods:** different clustering methods are selected for each network based on leaderboard performance or structural quality scores. |  |  |  |
| H1 | Either Louvain, Infomap, or a continuous optimization method (Li et al., 2016) are selected for each network. | 50 | R, F |
| H2 | Either Louvain, Infomap, SPICi, or DCut (Shao et al., 2016) are selected for each network. | 50 | W,R |
| H3 | Up to five different methods are applied to cluster networks, followed by filtering of modules based on structural quality metrics. | 40 | W,R, M, F |
| H4 | Up to nine different methods are applied in different combinations, followed by module filtering and post-processing steps. | 37 |  |
| H5 | Up to seven different methods are applied including an ensemble approach, followed by filtering and post-processing steps. | 31 | S,W,R,M,F |
| H6 | WGCNA followed by fast greedy community detection to refine modules. | 19 | R |
| H7 | No detailed description provided. | 14 | - |
| **Others** |  |  |  |
| O1 | Agglomerative algorithm that joins clusters based on the number of shared neighbors and the cluster sizes. | 36 | W,F |
| O2 | Two-way modules (dense bipartite subgraphs) are mined using a heuristic algorithm. | 33 | W,F |
| O3 | No detailed description provided. | 12 | - |

<sup>a</sup>Identifier (ID) of the method used throughout the paper.
<sup>b</sup>Overall score of the method as defined in **Fig. 2B**.
<sup>c</sup>Common pre- and post-processing steps. Pre-processing steps are coded as: (S) sparsification of networks and (W) rescaling of edge weights. Post-processing steps are coded as: (R) recursive break-down of large modules, (M) merging modules of invalid size followed by re-modularization, and (F) filtering modules according to a quality metric.

**Figure 2:**
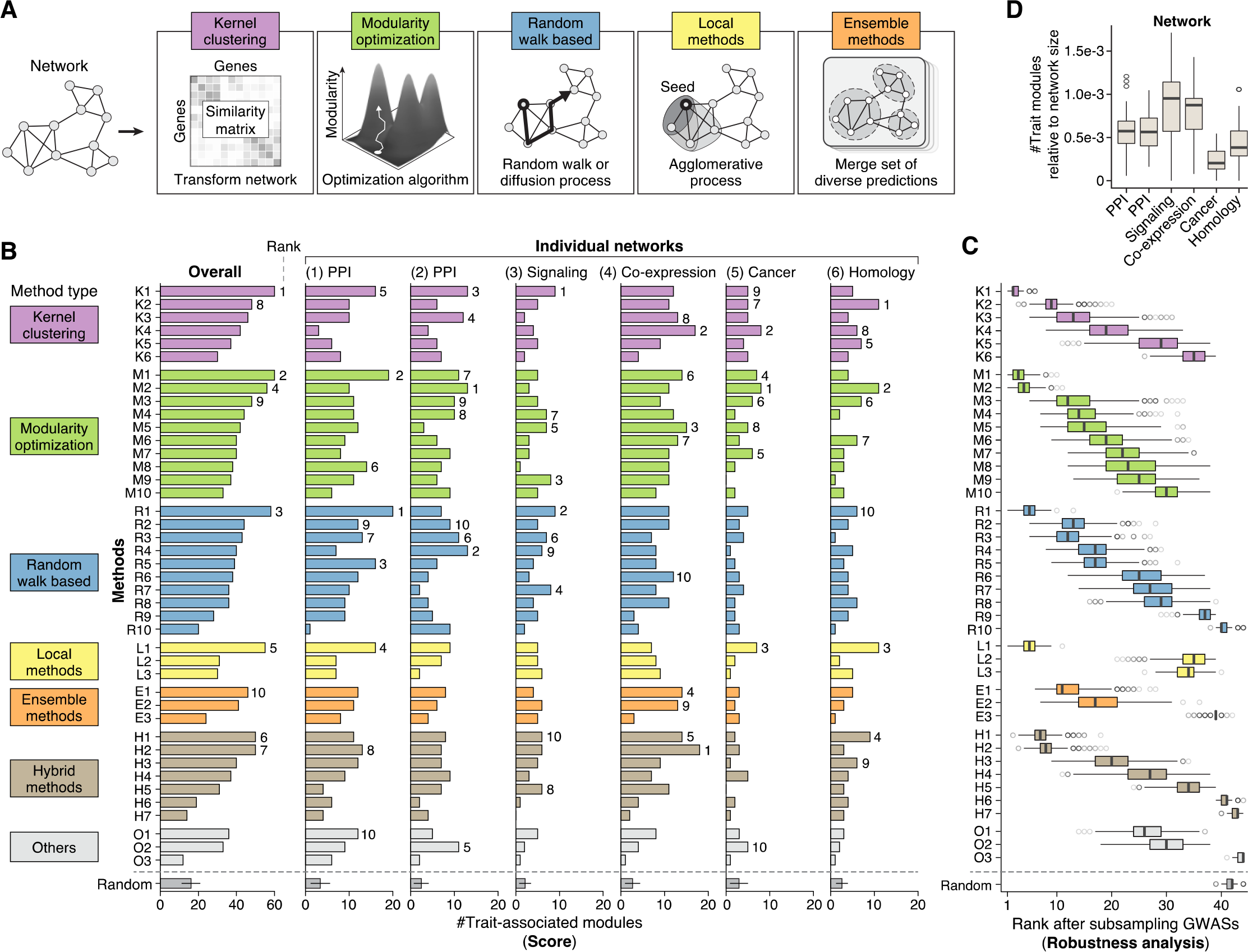
Assessment of module identification methods. **(A)** Main types of module identification approaches used in the challenge: kernel clustering methods transform and cluster the network adjacency matrix; modularity optimization methods rely on search algorithms to find modular decompositions that maximize a structural quality metric; random-walk-based methods take inspiration from diffusion processes over the network; local methods use agglomerative processes to grow modules from seed nodes; and ensemble methods merge alternative clusterings sampled either from stochastic runs of a given method or from a set of different methods. In addition, hybrid methods employ more than one of the above approaches and then pick the best modules according to a quality metric. See also Table 1. **(B)** Final scores of the 42 module identification methods applied in Sub-challenge 1 for each of the six networks, as well as the overall score summarizing performance across networks (same method identifiers as in Table 1). Scores correspond to the number of unique trait-associated modules identified by a given method in a network (evaluated using the hold-out GWAS set at 5% FDR, see Methods). Ranks are indicated for the top ten methods. The last row shows the performance of randomly generated modules. **(C)** Robustness of the overall ranking was evaluated by subsampling the GWAS set used for evaluation 1,000 times. For each method, the resulting distribution of ranks is shown as a boxplot. The rankings of method *K1* are substantially better than those of the remaining teams (Bayes factor < 3, see Methods). **(D)** Number of trait-associated modules per network. Boxplots show the number of trait-associated modules across methods, normalized by the size of the respective network. See also Fig. S1B.

### Top methods from different categories achieve comparable performance

In Sub-challenge 1, teams submitted a separate set of predicted modules for each of the six networks. We scored these predictions based on the number of trait-associated modules at 5% false discovery rate (FDR; Methods). The overall score used to rank methods in the challenge was defined as the total number of trait-associated modules across the six networks. (Module predictions, scoring scripts and full results are available in on the challenge website.)

The top five methods achieved comparable performance with scores between 55 and 60, while the remaining methods did not get to scores above 50 (Fig. 2B). To assess the robustness of the challenge ranking, we further scored all methods on 1,000 subsamples of the GWAS hold-out set (Methods). This analysis revealed a significant difference between the top-scoring method *K1* (method IDs are defined in Table 1) and the remaining methods (Fig. 2C). In addition, we repeated the scoring using four different FDR cutoffs: method *K1* ranked 1st in each case, while the performance of other methods varied (Fig. S1A). Moreover, method *K1* also obtained the top score in the leaderboard round. We conclude that although the final scores of the top 5 methods are close, method *K1* performed more robustly in diverse settings.

The top teams used different approaches: the best performers (*K1*) developed a novel kernel approach leveraging a diffusion-based distance metric (Cao et al., 2013, 2014) and spectral clustering (Ng et al., 2001); the runner-up team (*M1*) extended different modularity optimization methods with a resistance parameter that controls the granularity of modules (Arenas et al., 2008); and the third-ranking team (*R1*) used a random-walk method based on multi-level Markov clustering with locally adaptive granularity to balance module sizes (Satuluri et al., 2010). Interestingly, teams employing the widely-used Weighted Gene Co-expression Network Analysis (WGCNA) tool (Langfelder and Horvath, 2008), which relies on hierarchical clustering to detect modules, did not perform competitively in this challenge (rank 35, 37 and 41).

Four different method categories are represented among the top five performers, suggesting that no single approach is inherently superior for module identification in molecular networks. Rather, performance depends on the specifics of each individual method, including the strategy used to define the resolution of the modular decomposition (the number and size of modules, discussed more in detail in Section “Complementarity of different module identification approaches”). Notably, the two runner-up teams (*M1* and *R1*) both used methods specifically designed to control the resolution of modules. Pre-processing steps also affected performance: many of the top teams first sparsified the networks by discarding weak edges. A notable exception is the top method (*K1*), which performed robustly without any pre-processing of the networks.

The challenge also allows us to explore how informative different types of molecular networks are for finding modules underlying complex traits. In absolute numbers, methods recovered the most trait-associated modules in the co-expression and protein-protein interaction networks (Fig. S1B). However, relative to the network size, the signaling network contained the most trait-associated modules (Fig. 2D). These results are consistent with the importance of signaling pathways for many of the considered traits and diseases. The cancer cell line and homology-based networks, on the other hand, were less relevant for the traits in our GWAS compendium and thus comprised only few trait-associated modules.

### Complementarity of different module identification approaches

We next asked whether predictions from different methods and networks tend to capture the same or complementary modules. To this end, we developed a pairwise similarity metric for module predictions, which we applied to the complete set of 252 module predictions from Sub-challenge 1 (42 methods × 6 networks, Methods). We find that similarity of module predictions is primarily driven by the underlying network and not the method category (Fig. 3A). When comparing module predictions between methods, we find that top-performing methods produced dissimilar clusterings, suggesting that they capture complementary functional modules (Fig. S3A).

**Figure 3:**
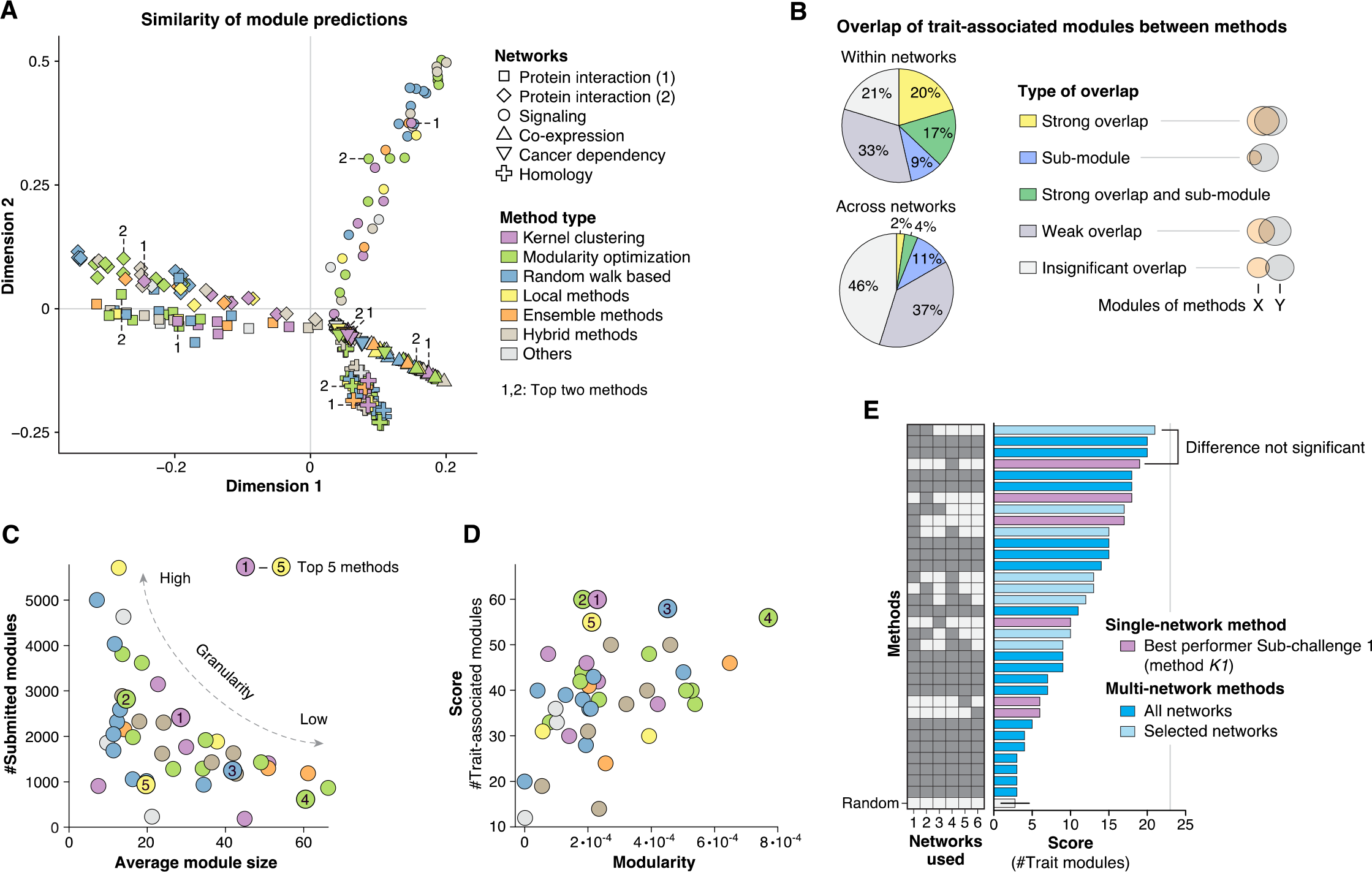
Complementarity of module predictions from different methods and networks. **(A)** Similarity of module predictions from different methods (color) and networks (shape). The closer two points are in the plot, the more similar are the corresponding module predictions (multidimensional scaling, see Methods). Top performing methods tend to be located far from the origin (the top two methods are highlighted for each network). Top methods do not cluster close together, suggesting dissimilar modular decompositions (see also Fig. S3A). **(B)** Comparison of GWAS trait-associated modules identified by all challenge methods. Pie-charts show the percentage of trait modules that show overlap with at least one trait module from a different method in the same network (top) and in different networks (bottom). We distinguish between strong overlap, sub-modules, weak but significant overlap, and insignificant overlap (Methods). **(C)** Total number of predicted modules versus average module size for each method (same color scheme as in Panel A). There is a roughly inverse relationship between module number and size because modules had to be non-overlapping and did not have to cover all genes. The top five methods (highlighted) produced modular decompositions of varying granularity. See also Figs. S3B-E. **(D)** Challenge score (number of trait-associated modules) versus modularity is shown for each method (same color scheme as in Panel A). Modularity is a topological quality metric for modules based on the fraction of within-module edges (Newman and Girvan, 2004). While there is modest correlation between the two metrics (r=0.45), the methods with the highest challenge score are not necessarily those with the highest modularity, presumably because the intrinsic scale of modularity is not optimal for the task considered in the challenge. **(E)** Final scores of multi-network module identification methods in Sub-challenge 2 (evaluated using the hold-out GWAS set at 5% FDR, see Methods). For comparison, the overall best-performing method from Sub-challenge 1 is also shown (method *K1*, purple). Teams used different combinations of the six challenge networks for their multi-network predictions (shown on the left): the top-performing team relied exclusively on the two protein-protein interaction networks. The difference between the top single-network module predictions and the top multi-network module predictions is not significant when sub-sampling the GWASs (Fig. S1D). The last row shows the performance of randomly generated modules

These observations can be confirmed by evaluating the overlap between trait-associated modules from different methods. Within the same network, only 46% of trait modules are recovered by multiple methods with good agreement (high overlap or submodules, Fig. 3B). Across different networks, the number of recovered modules with substantial overlap is even lower (17%). Thus, the majority of trait modules are method- and network-specific. This suggests that users should not rely on a single method or network to find trait-relevant modules.

The modules produced by different methods also vary in terms of their structural properties. For example, submissions included between 16 and 1552 modules per network, with an average module size ranging from 7 to 66 genes. Consistent with an analysis of potential scoring biases that we performed prior to the challenge, neither the number nor the size of submitted modules correlates with performance (Figs. 3C, **S3B-E**). The scoring metric is thus not biased towards a specific module granularity, rather different methods captured trait-relevant modules at varying levels of granularity. Topological quality metrics of modules such as modularity showed only modest correlation with the challenge score (Fig. 3D), highlighting the need to empirically assess module identification methods for a given task.

### Multi-network module identification methods did not provide added power

In Sub-challenge 2, teams submitted a single modularization of the genes, for which they could leverage information from all six networks together. While some teams developed dedicated multi-network (multi-layer) community detection methods (De Domenico et al., 2015; Didier et al., 2015), the majority of teams first merged the networks in some way and then applied single-network methods.

It turned out to be very difficult to effectively leverage complementary networks for module identification. While three teams achieved marginally higher scores than single-network module predictions, the difference is not significant (Figs. 3E, **S1C**). Moreover, the best-scoring team simply merged the two protein interaction networks (the two most similar networks, Fig. S2E), discarding the other types of networks. Since no significant improvement over single-network methods was achieved, the winning position of Sub-challenge 2 was declared vacant.

### Integration of challenge submissions leads to robust consensus modules

Integration of multiple team submissions often leads to robust consensus predictions in crowdsourced challenges (Marbach et al., 2012). We therefore developed an ensemble approach to derive consensus modules from team submissions. To this end, module predictions from different methods were integrated in a consensus matrix *C*, where each element *c*_*ij*_ is proportional to the number of methods that put gene *i* and *j* together in the same module. The consensus matrix was then clustered using the top-performing module identification method from the challenge (Fig. S2A, Methods).

We generated consensus modules for each challenge network by applying this approach to the top 21 (50%) of methods from the leaderboard round. The score of the consensus modules outperforms the top individual method predictions in both sub-challenges, highlighting the value of our community-based module collection (Fig. S1). However, when applied to fewer methods, the performance of the consensus drops (Fig. S2C), suggesting that further work is needed to make our consensus approach practical outside of a challenge context.

### Network modules reveal trait-specific and shared pathways

We next sought to explore biological properties of trait-associated modules discovered by the challenge participants. In what follows, we focus on the single-network predictions from Sub-challenge 1. The most trait-associated modules were found for immune-related, psychiatric, blood cholesterol and anthropometric traits, for which high-powered GWAS are available that are known to show strong pathway enrichment (Fig. 4A).

**Fig. 4:**
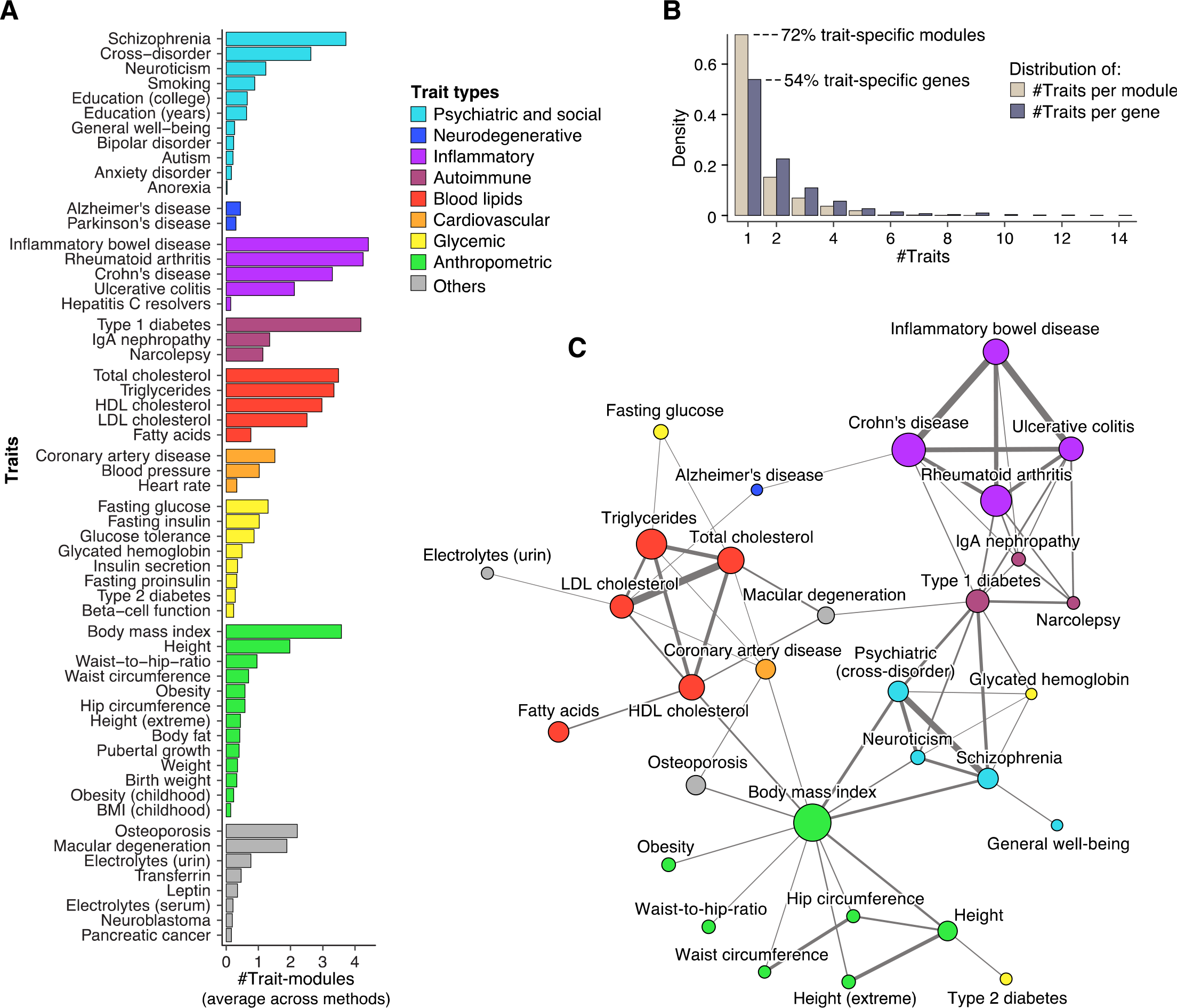
Overlap between modules associated with different traits and diseases. **(A)** Average number of trait-associated modules identified by challenge methods for each trait. For traits where multiple GWASs were available, results for the best-powered study are shown. **(B)** Histograms showing the number of distinct traits per trait-associated module (brown) and gene (grey). 72% of trait-associated modules are specific to a single trait, while the remaining 28% are hits for multiple traits. In contrast, only 54% of trait-associated genes are specific to a single trait. **(C)** Trait network showing similarity between GWAS traits based on overlap of associated modules (force-directed graph layout). Node size corresponds to the number of genes in trait-associated modules and edge width corresponds to the degree of overlap (Jaccard index; only edges for which the overlap is significant are shown, see Methods). Traits without any edges are not shown. Traits of the same type (color) tend to cluster together, indicating shared pathways.

Significant GWAS loci often show association to multiple traits. Across our GWAS compendium, we found that 46% of trait-associated genes but only 28% of trait-associated modules are associated with multiple traits (Fig. 4B). Thus, mapping genes onto network modules may help disentangling trait-specific pathways at shared loci.

We further asked which traits are similar in terms of the implicated network components. To this end, we considered the union of all genes within network modules associated with a given trait (called “trait-module genes”). We then evaluated the pairwise similarity of traits based on the significance of the overlap between the respective trait-module genes (Methods). Trait relationships thus inferred are consistent with known biology and comorbidities between the considered traits and diseases (Fig. 4C). For example, consistent with its pathophysiological basis, age-related macular degeneration shares network components with cholesterol and immune traits, while coronary artery disease shows similarity with established risk factors (cholesterol levels, body mass index) and osteoporosis, which is epidemiologically and biologically linked (atherosclerotic calcification and bone mineralization involve related pathways).

### Trait-associated modules implicate core disease genes and pathways

Trait-associated modules typically include many genes that do not show any signal in the respective GWAS. A key question is whether modules correctly predict such genes as being relevant for that trait or disease. We first consider a module from the consensus analysis that shows association to height -- a classic polygenic trait -- as an example. In the GWAS that was used to identify this module there are only three module genes that show association to height, while the remaining genes are predicted to play a role in height solely because they are members of this module (Fig. 5A). We sought to evaluate such candidate genes for height as well as other traits using higher-powered GWASs, ExomeChip data, monogenic disease genes and functional annotations.

**Figure 5:**
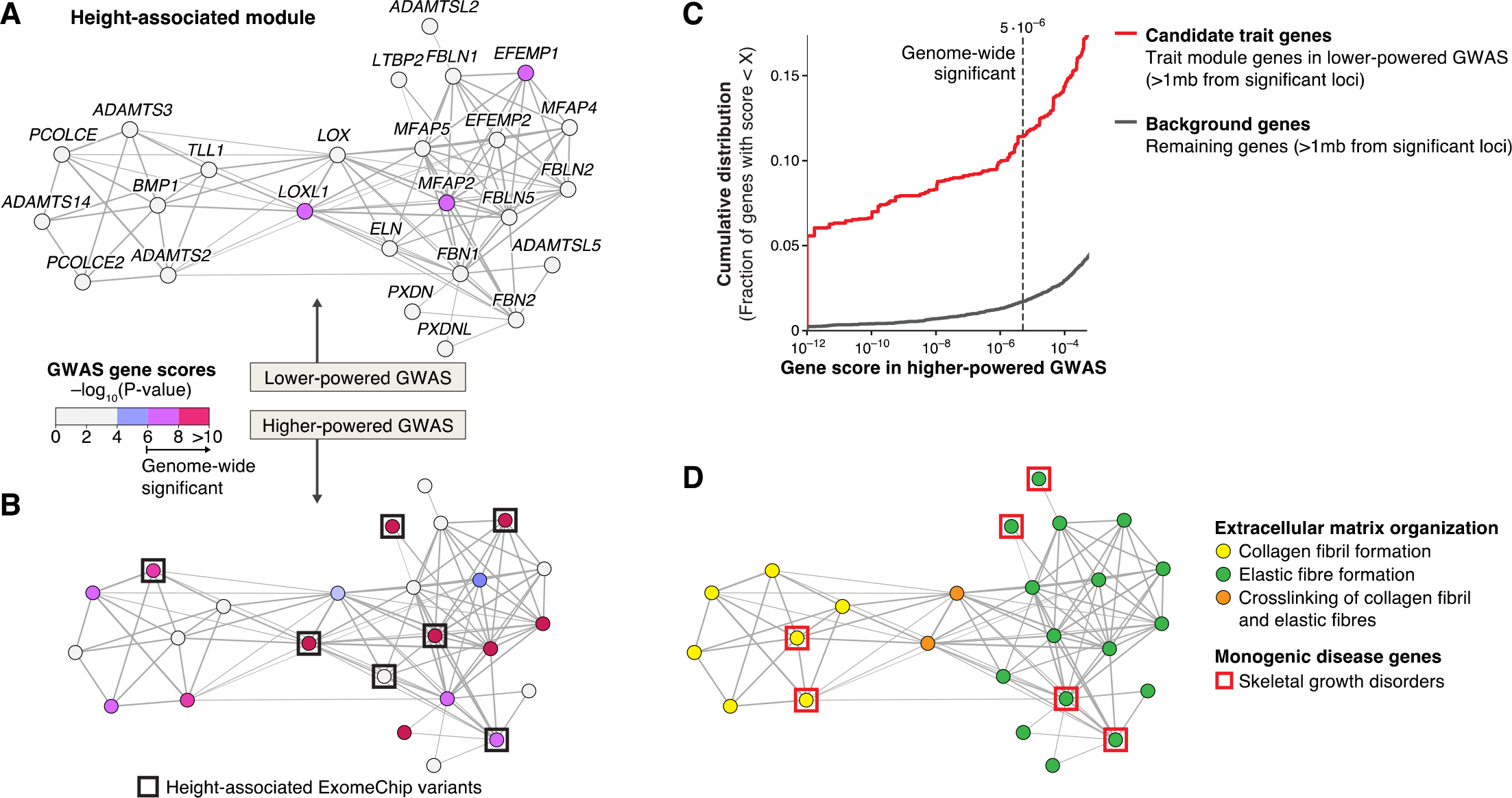
Support of trait-module genes in diverse datasets. **(A)** Example module of the consensus analysis in the STRING protein interaction network (force-directed graph layout). The module shows modest association to height (*q*-value = 0.04) in the GWAS by Randall et al. (2013) (lower-powered than the GWAS shown in Panel B). Color indicates GWAS gene scores. The signal is driven by three genes from different loci with significant scores (pink), while the remaining genes (grey) are predicted to be involved in height because of their module membership. **(B)** The module from **Panel A** is supported in the higher-powered GWAS (Wood et al., 2014) (*q*-value = 0.005). 45% of candidate trait genes (grey in **Panel A**) are confirmed (pink). In addition, 28% of module genes have coding variants associated to height in an independent ExomeChip study published after the challenge (Marouli et al., 2017) (black squares, enrichment *p*-value = 1.9E-6 [one-sided Fisher’s exact test]). See also Fig. S4B. **(C)** Support of candidate trait genes across eight different traits for which lower- and higher-powered GWASs are available in our hold-out set. The lower-powered GWASs were used to predict candidate trait genes, i.e., genes within trait modules that do not show any signal (GWAS gene score <4) and that are located far away (>1mb) from any significant GWAS locus (cf. grey genes in **Panel A**). The plot shows the cumulative distribution of gene scores in the higher-powered GWASs for candidate trait genes (red line) and all other genes (grey line, see Methods). **(D)** Functional annotation of genes in the height-associated module from **Panel A**. Genes implicated in monogenic skeletal growth disorders are highlighted (red squares, enrichment *p*-value = 7.5E-4). See also Table S2.

There are eight traits for which we have both an older (lower-powered) and more recent (higher-powered) GWAS in our hold-out set: height, schizophrenia, ulcerative colitis, Crohn’s disease, rheumatoid arthritis, and three blood lipid traits (Fig. S4A). We can thus identify trait modules and candidate genes using the lower-powered GWAS and then evaluate how well they are supported in higher-powered GWAS (a common approach used to assess methods for GWAS gene prioritization, see Methods). Indeed, while only 3 genes in the height module introduced above are associated to height in the lower-powered GWAS (Randall et al., 2013), 13 module genes are confirmed in the higher-powered GWAS (Wood et al., 2014) and 6 module genes further comprise coding variants associated to height in an independent ExomeChip study (Marouli et al., 2017) (Fig 5B). Similar results are obtained when evaluating module predictions from all challenge methods across the eight above-mentioned traits: a substantial fraction of module genes that do not show any signal and are located far from any significant locus in the lower-powered GWAS are subsequently confirmed by the higher-powered GWAS (Fig. 5C). This demonstrates that modules are predictive for trait-associated genes and could thus be used to prioritize candidate genes for follow-up studies, for instance.

We next explored the biological function and clinical relevance of identified trait modules. For example, the height module discussed above consists of two submodules comprising extracellular matrix proteins responsible for, respectively, collagen fibril and elastic fiber formation -- pathways that are essential for growth (Fig. 5D). Indeed, mutations of homologous genes in mouse lead to abnormal elastic fiber morphology (Table S2) and one out of four module genes are known to cause monogenic skeletal growth disorders in human (Fig. 5D). For example, the module gene *BMP1* (*Bone Morphogenic Protein 1*) causes osteogenesis imperfecta, which is associated with short stature. Interestingly, *BMP1* does not show association to height in current GWAS and ExomeChip studies (Fig. 5A,B), demonstrating how network modules can implicate additional disease-relevant pathway genes (see Fig. S4B for a systematic comparison of trait modules with independent disease gene sets from the literature).

To evaluate more generally whether trait-associated modules correspond to generic or disease-specific pathways, we visualized and tested modules for functional enrichment of Gene Ontology (GO) annotations, mouse mutant phenotypes, and diverse pathway databases. In order to account for annotation bias of well-studied genes (Glass and Girvan, 2014), we employed a non-central hypergeometric test (Methods). We find that the majority of trait modules reflect core disease-specific pathways. For example, in the first protein-protein interaction network only 33% of trait modules from the consensus analysis have generic functions, such as epigenetic gene silencing for modules associated with schizophrenia and body mass index; the remaining 66% of trait modules correspond to core disease-specific pathways, some of which are therapeutic targets (Fig. 6 and Tables S3, **S4**). Examples include a module associated with rheumatoid arthritis that comprises the B7:CD28 costimulatory pathway required for T cell activation, which is blocked by an approved drug (Fig. 6A); a module associated with inflammatory bowel disease corresponding to cytokine signalling pathways mediated by Janus kinases (JAKs), which are therapeutically being targeted at multiple levels (Fig. 6B); and a module associated with myocardial infarction that includes the NO/cGMP signaling cascade, which plays a key role in cardiovascular pathophysiology and therapeutics (Fig. 6C). We further applied our pipeline to a GWAS on IgA nephropathy (IgAN) obtained after the challenge, a disease with poorly understood etiology and no effective therapy (Kiryluk et al., 2014). IgAN is an autoimmune disorder that manifests itself by deposition of immune complexes in the kidney’s glomeruli, triggering inflammation (glomerulonephritis) and tissue damage. The best-performing challenge method (*K1*) revealed one IgAN-specific module. The module implicates complement and coagulation cascades, pointing to the chemokine *PF4V1* as a novel candidate gene (Fig. 6D). In support of the function of this module in IgAN, top enriched mouse mutant phenotypes for module gene homologs are precisely “glomerulonephritis” and “abnormal blood coagulation” (Fig. S5).

**Figure 6:**
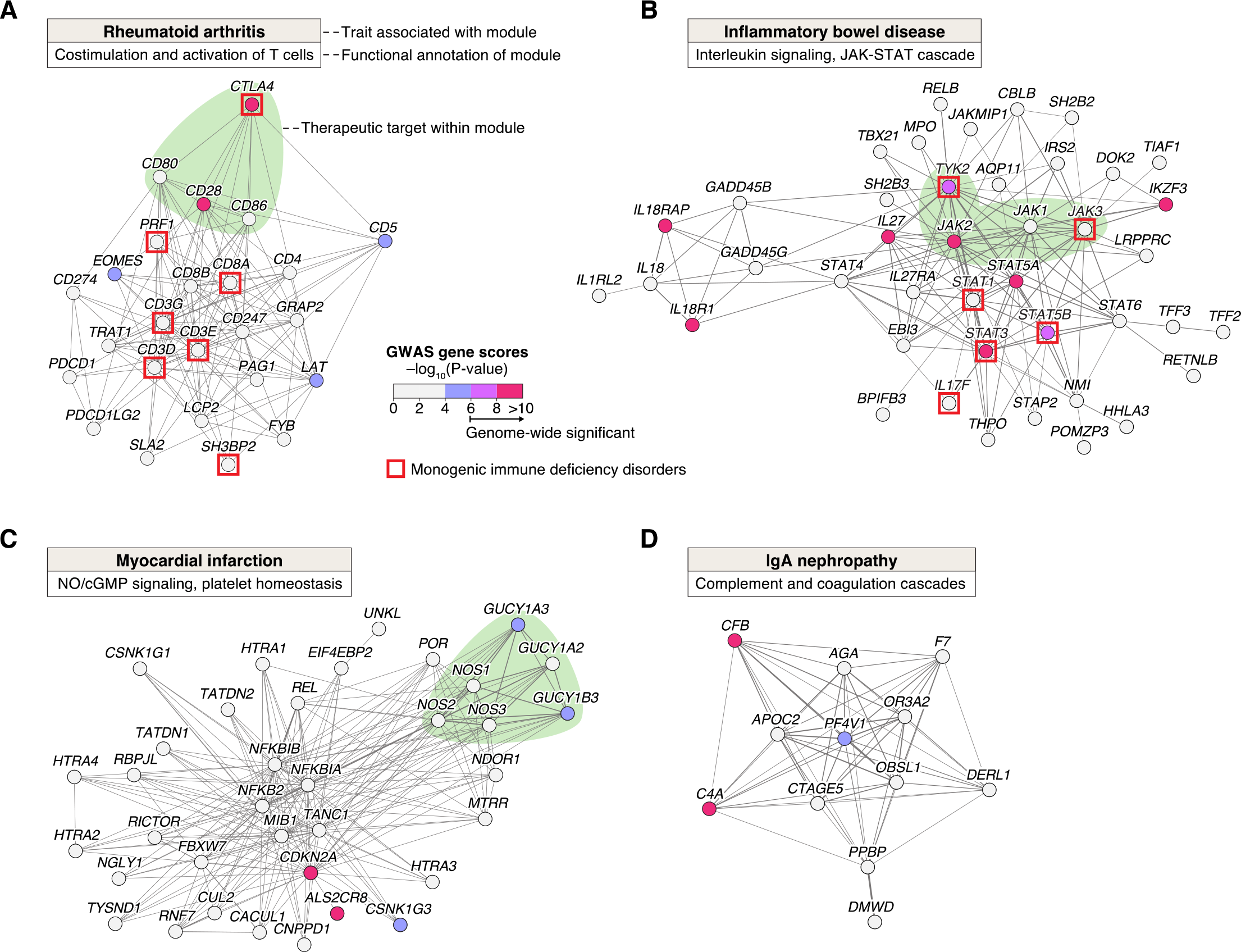
Example trait modules comprising therapeutically relevant pathways. **(A, B and C)** Three trait-associated modules in the STRING protein interaction network from the consensus analysis (similar results were obtained for other modules and traits, Tables S3, **S4**). Node colors correspond to gene scores in the respective GWAS. For the two inflammatory disorders (A and B), red squares indicate genes causing monogenic immunodeficiency disorders (enrichment *p*-values of 4.1E-8 and 1.2E-6, respectively [one-sided Fisher’s exact test]). **(A)** Module associated with rheumatoid arthritis (*q*-value = 0.04) involved in T cell activation. A costimulatory pathway is highlighted green: T cell response is regulated by activating (*CD28*) and inhibitory (*CTLA4*) surface receptors, which bind B7 family ligands (*CD80* and *CD86*) expressed on the surface of activated antigen-presenting cells. The therapeutic agent CTLA4-Ig binds and blocks B7 ligands, thus inhibiting T cell response. **(B)** A cytokine signalling module associated with inflammatory bowel disease (*q*-value = 0.0006). The module includes the four known Janus kinases (*JAK1-3* and *TYK2*, highlighted green), which are engaged by cytokine receptors to mediate activation of specific transcription factors (*STATs*). Inhibitors of JAK-STAT signaling are being tested in clinical trials for both ulcerative colitis and Crohn’s disease (Neurath, 2017). **(C)** Module associated with myocardial infarction (*q*-value = 0.0001). The module includes two main components of the NO/cGMP signaling pathway (highlighted green): endothelial nitric oxide synthases (*NOS1-3*), which produce the gas nitric oxide (NO) used as signal transmitter, and soluble guanylate cyclases (*GUCY1A2*, *GUCY1A3* and *GUCY1B3*), which sense NO leading to formation of cGMP. The cGMP signal inhibits platelet aggregation and leads to vascular smooth muscle cell relaxation; it is a therapeutic target for cardiovascular disease as well as erectile dysfunction (Kraehling and Sessa, 2017). **(D)** Module associated to IgA nephropathy (IgAN; *q*-value = 0.04). The module was identified using the best-performing method (*K1*) in the InWeb protein interaction network. Besides finding complement factors that are known to play a role in the disease (*CFB* and *C4A*), the module implicates novel candidate genes such as the chemokine *Platelet Factor 4 Variant 1* (*PF4V1*) from a sub-threshold locus, and is enriched for coagulation cascade, a process known to be involved in kidney disease (Madhusudhan et al., 2016) (see also Fig. S5).

## Discussion

Large-scale network data are becoming pervasive in many areas ranging from the digital economy to the life sciences. While analysis goals vary across fields, unsupervised detection of network communities remains an essential task in many applications of interest. We have conducted a critical assessment of module identification methods on real-world networks, providing much-needed guidance for users. The community-based challenge enabled comprehensive and impartial assessment, avoiding the “self-assessment trap” that leads researchers to consciously or unconsciously overestimate performance when evaluating their own algorithms (Norel et al., 2011). While it is important to keep in mind that the exact ranking of methods -- as in any benchmark -- is specific to the task and datasets considered, we believe that the resulting collection of top-performing module identification tools and methodological insights will be broadly useful for modular analysis of complex networks in biology and other domains.

In addition to providing a cross section of established approaches, the collection of contributed methods also includes novel algorithms that further advance the state-of-the-art (notably, the best-performing method). Kernel clustering, modularity optimization, random-walk-based and local methods were all represented among the top performers, suggesting that no single type of approach is inherently superior. In contrast, some widely used methods for gene network analysis, including WGCNA and hierarchical clustering, did not perform competitively. Moreover, while most published studies in network biology rely on a single clustering method, the results of this challenge demonstrate the value of applying multiple methods from different categories, which favors the detection of complementary types of modules. We find that the top four challenge methods (*K1*, *M1*, *R1* and *M2*) already offer substantial diversity (Fig. S3F). It should be noted that the larger number of modules also results in a higher multiple testing burden in any subsequent analyses (e.g., functional enrichment testing) and that modules from different methods may overlap. When a single non-overlapping partition is needed, the best-performing challenge method (*K1*) is a good choice as it functioned robustly in diverse settings.

The challenge also emphasized the importance of the resolution (size and number of modules), which critically affected results. Biological networks typically have a hierarchical modular structure, which implies that disease-relevant pathways can be captured at different levels (Ravasz et al., 2002). We have shown that our scoring method is not biased towards a specific module size, rather the optimal resolution is method- and network-specific (Fig. S3B-E). Top-performing challenge methods allowed the resolution to be tuned. Although setting the “right” resolution can be challenging for users, this critical point should not be sidestepped. We recommend that users experiment with different resolutions and use the settings optimized by teams for the different types of networks as guidance.

Our analysis showed that signaling, protein-protein interaction and co-expression networks comprise complementary trait-relevant modules, and that the choice of network has a stronger effect than the choice of method on the resulting modules (Fig. 3A,B). Further improving the quality of available network datasets is thus crucial and considering different types of networks is clearly advantageous. However, multi-network module identification methods that attempted to reveal integrated modules across these networks failed to significantly improve predictions compared to methods that considered each network individually. This is a sobering result for the emerging field of multi-network methods, challenging the common assumption that integration of complementary types of networks allows more accurate identification of functional modules. Possibly, the networks of the challenge were not sufficiently related – multi-network methods may perform better on networks from the same tissue- and disease-context (Krishnan et al., 2016). An avenue for future work is thus to explore how network relatedness affects performance of multi-network module identification.

Here we assessed unsupervised clustering methods that define modules based solely on the topological structure of networks, unbiased by existing biological knowledge. Additional biological information, such as GWAS data and functional annotations, was integrated afterwards to characterize the predicted modules. In contrast, supervised or integrative network analysis methods rely on additional biological data from the start, e.g., by projecting known disease genes onto the network and finding modules that span these genes (Cowen et al., 2017). Unsupervised and supervised approaches have different strengths and limitations, the former being generally preferred for unbiased identification of network modules whereas the latter enable identification of context-specific modules, for example. While our challenge resources may also provide a foundation for assessment of certain types of supervised or integrative clustering approaches, this is outside the scope of the present study. Likewise, methods for detection of overlapping modules (Ihmels et al., 2002) may also be assessed using the challenge framework, but will require development of novel scoring metrics that take module overlap into account.

The collective effort of over 400 challenge participants resulted in a unique compendium of modules for the different types of molecular networks considered. By leveraging the “wisdom of crowds” we generated robust consensus modules. While most modules partly reflect known pathways or functional gene categories, which they reorganize and expand with additional genes, other modules may correspond to yet uncharacterized pathways. The consensus modules (gene sets) thus constitute a novel data-driven pathway collection, which may complement existing pathway collections in a range of applications (e.g., for interpretation of gene expression data using gene set enrichment analysis).

There is continuing debate over the value of GWASs for revealing disease mechanisms and therapeutic targets. Indeed, the number of GWAS hits continues to grow as sample sizes increase, but the bulk of these hits may not correspond to core genes with specific roles in disease etiology. An “omnigenic” model recently proposed by Boyle et al. (2017) explains this observation by the high interconnectivity of molecular networks, which implies that most of the expressed genes in a disease-relevant tissue are likely to be at least weakly connected to core genes and may thus have non-zero effects on that disease. Indeed, disease-associated genes tend to coalesce in regulatory networks of tissues that are specific to that disease (Marbach et al., 2016). While thousands of genes may show association to a given disease, we have demonstrated that specific disease modules comprising only dozens of genes can be identified within networks. These modules are more disease-specific than individual genes, reveal pathway-level similarity between diseases, accurately prioritize candidate genes, and correspond to core disease pathways in the majority of cases. This is consistent with the omnigenic model and the robustness of biological networks: presumably, the many genes that influence disease indirectly are broadly distributed across network modules, while core disease genes cluster in specific pathways underlying pathophysiological processes (Pers et al., 2015; Sullivan and Posthuma, 2015).

In this study we used generic networks, not context-specific networks, because the focus was on method assessment across diverse disorders. In the near future, we expect much more detailed maps of tissue- and disease-specific networks, along with diverse high-powered genetic datasets, to become available. We hope that the challenge resources will be instrumental in dissecting these networks and will provide a solid foundation for developing integrative methods to reveal the cell types and causal circuits implicated in human disease.

## Consortia

The contributing members of the DREAM Module Identification Challenge Consortium are: Fabian Aicheler,^1^ Nicola Amoroso,^2,3^ Alex Arenas,^4^ Karthik Azhagesan,^5-,7^ Aaron Baker,^8-10^ Michael Banf,^11^ Serafim Batzoglou,^12^ Anaïs Baudot,^13^ Roberto Bellotti,^2,3,14^ Sven Bergmann,^15,16^ Keith A. Boroevich,^17^ Christine Brun,^18-19^ Stanley Cai,^20,93,94^ Michael Caldera,^21^ Alberto Calderone,^22^ Gianni Cesareni,^22^ Weiqi Chen,^23^ Christine Chichester,^24^ Sarvenaz Choobdar,^15-16^ Lenore Cowen,^25-26^ Jake Crawford,^25^ Hongzhu Cui,^27^ Phuong Dao,^46^ Manlio De Domenico,^4,29^ Andi Dhroso,^27^ Gilles Didier,^13^ Mathew Divine,^1^ Antonio del Sol,^36^ Tao Fang,^96^ Xuyang Feng,^30^ Jose C. Flores-Canales,^31-32^ Santo Fortunato,^33^ Anthony Gitter,^8,9,10^ Anna Gorska,^34^ Yuanfang Guan,^35^ Alain Guénoche,^13^ Sergio Gómez,^4^ Hatem Hamza,^24^ András Hartmann,^36^ Shan He,^23^ Anton Heijs,^37^ Julian Heinrich,^1^ Benjamin Hescott,^38^ Xiaozhe Hu,^26^ Ying Hu,^39^ Xiaoqing Huang,^46^ V. Keith Hughitt,^40-41^ Minji Jeon,^42^ Lucas Jeub,^33^ Nathan Johnson,^27^ Keehyoung Joo,^32,43^ InSuk Joung,^31-32^ Sascha Jung,^36^ Susana G. Kalko,^36^ Piotr J. Kamola,^17^ Jaewoo Kang,^42,44^ Benjapun Kaveelerdpotjana,^23^ Minjun Kim,^45^ Yoo-Ah Kim,^46^ Oliver Kohlbacher,^1,47-48^ Dmitry Korkin,^27,49-50^ Kiryluk Krzysztof,^51^ Khalid Kunji,^52^ Zoltàn Kutalik,^16,53^ Kasper Lage,^54-56^ David Lamparter,^15-16,57^ Sean Lang-Brown,^58^ Thuc Duy Le,^59-60^ Jooyoung Lee,^31-32^ Sunwon Lee,^42^ Juyong Lee,^61^ Dong Li,^23^ Jiuyong Li,^60^ Junyuan Lin,^26^ Lin Liu,^60^ Antonis Loizou,^62^ Zhenhua Luo,^63^ Artem Lysenko,^17^ Tianle Ma,^64^ Raghvendra Mall,^52^ Daniel Marbach,^15-16,96^ Tomasoni Mattia,^15-16^ Mario Medvedovic,^65^ Jörg Menche,^21^ Johnathan Mercer,^54,56^ Elisa Micarelli,^22^ Alfonso Monaco,^3^ Felix Müller,^21^ Rajiv Narayan,^66^ Oleksandr Narykov,^50^ Ted Natoli,^66^ Thea Norman,^67^ Sungjoon Park,^42^ Livia Perfetto,^22^ Dimitri Perrin,^68^ Stefano Pirrò,^22^ Teresa M. Przytycka,^46^ Xiaoning Qian,^69^ Karthik Raman,^5-7^ Daniele Ramazzotti,^12^ Emilie Ramsahai,^95^ Balaraman Ravindran,^70,6,7^ Philip Rennert,^71^ Julio Saez-Rodriguez,^7-,73^ Charlotta Schärfe,^1^ Roded Sharan,^74^ Ning Shi,^23^ Wonho Shin,^44^ Hai Shu,^75^ Himanshu Sinha,^5,6,7^ Donna K. Slonim,^25^ Lionel Spinelli,^18^ Suhas Srinivasan,^49^ Aravind Subramanian,^66^ Christine Suver,^76^ Damian Szklarczyk,^77^ Sabina Tangaro,^3^ Suresh Thiagarajan,^78^ Laurent Tichit,^13^ Thorsten Tiede,^1^ Beethika Tripathi,^70,6,7^ Aviad Tsherniak,^66^ Tatsuhiko Tsunoda,^17,79,80^ Dénes Türei,^72^ Ehsan Ullah,^52^ Golnaz Vahedi,^20,93,94^ Alberto Valdeolivas,^13,82^ Jayaswal Vivek,^83^ Christian von Mering,^77^ Andra Waagmeester,^37^ Bo Wang,^12^ Yijie Wang,^46^ Barbara A. Weir,^84-85^ Shana White,^65^ Sebastian Winkler,^1^ Ke Xu,^86^ Taosheng Xu,^87^ Chunhua Yan,^39^ Liuqing Yang,^88^ Kaixian Yu,^75^ Xiangtian Yu,^89^ Gaia Zaffaroni,^36^ Mikhail Zaslavskiy,^90^ Tao Zeng,^89^ Jitao D. Zhang,^96^ Lu Zhang,^12^ Weijia Zhang,^60^ Lixia Zhang,^65^ Xinyu Zhang,^86^ Junpeng Zhang,^91^ Xin Zhou,^12^ Jiarui Zhou,^23^ Hongtu Zhu,^75^ Junjie Zhu,^92^ Guido Zuccon.^68^

^1^Applied Bioinformatics, Center for Bioinformatics, University of Tuebingen, Sand 14, 72076 Tuebingen, Germany. ^2^Department of Physics ’Michelangelo Merlin’, University of Bari ’Aldo Moro’, Via G. Amendola 173, 70126 Bari, Italy. ^3^INFN, Sezione di Bari, Via A. Orabona 4, 70125 Bari, Italy. ^4^Departament d’Enginyeria Informàtica i Matemàtiques, Universitat Rovira i Virgili, Tarragona, Spain. ^5^Department of Biotechnology, Bhupat and Jyoti Mehta School of Biosciences, Indian Institute of Technology Madras, Chennai, India. ^6^Initiative for Biological Systems Engineering (IBSE), Indian Institute of Technology Madras. ^7^Robert Bosch Centre for Data Science and Artificial Intelligence (RBC-DSAI), Indian Institute of Technology Madras. ^8^Department of Biostatistics and Medical Informatics, University of Wisconsin-Madison, Madison, Wisconsin, USA. ^9^Department of Computer Sciences, University of Wisconsin-Madison, Madison, Wisconsin, USA. ^10^Morgridge Institute for Research, Madison, Wisconsin, USA. ^11^Department of Plant Biology, Carnegie Institution for Science, Stanford, USA. ^12^Department of Computer Science, Stanford University, USA. ^13^Aix Marseille Univ, CNRS, Centrale Marseille, I2M, UMR 7373, Marseille, France. ^14^Centro TIRES, Via G. Amendola 173, 70126 Bari, Italy. ^15^Department of Computational Biology, University of Lausanne, Lausanne, Switzerland. ^16^Swiss Institute of Bioinformatics, Lausanne, Switzerland. ^17^RIKEN Center for Integrative Medical Sciences, Yokohama, Japan. ^18^Aix Marseille Univ, INSERM, TAGC, UMR1090, Marseille, France. ^19^CNRS, Marseille, France. ^20^Department of Genetics, Perelman School of Medicine at the University of Pennsylvania, Philadelphia, Pennsylvania, USA. ^21^CeMM Research Center for Molecular Medicine of the Austrian Academy of Sciences, Vienna, Austria. ^22^Bioinformatics and Computational Biology Unit, Department of Biology, Tor Vergata University, Italy. ^23^School of Computer Science, The University of Birmingham, Birmingham, UK. ^24^Nestle Institute of Health Sciences, Lausanne, Switzerland.^25^Department of Computer Science, Tufts University, Medford, MA, USA. ^26^Department of Mathematics, Tufts University, Medford, MA, USA. ^27^Bioinformatics and Computational Biology Program, Worcester Polytechnic Institute, Worcester, MA, USA. ^29^Fondazione Bruno Kessler, Via Sommarive 18, 38123 Povo, Italy. ^30^Department of Cancer Biology, University of Cincinnati, Cincinnati, OH, USA. ^31^Center for In Silico Protein Science, Korea Institute for Advanced Study, Seoul, Korea. ^32^School of Computational Sciences, Korea Institute for Advanced Study, Seoul, Korea. ^33^School of Informatics, Computing and Engineering, Indiana University, Bloomington, USA. ^34^Algorithms in Bioinformatics, Center for Bioinformatics, University of Tuebingen, Sand 14, 72076 Tuebingen, Germany. ^35^Department of Computational Medicine and Bioinformatics, University of Michigan, Ann Arbor, MI, 48109. ^36^LCSB - Luxembourg Centre for Systems Biomedicine, University of Luxembourg, Esch-sur-Alzette, Luxembourg. ^37^Micelio, 2180 Antwerp, Belgium. ^38^College of Computer and Information Science, Northeastern University, Boston, MA, USA. ^39^National Cancer Institute, Center for Biomedical Informatics & Information Technology, 9609 Medical Center Drive, Bethesda, MD 20850, USA. ^40^Center for Bioinformatics and Computational Biology, University of Maryland, College Park, Maryland, USA. ^41^Department of Cell Biology and Molecular Genetics, University of Maryland, College Park, Maryland, USA. ^42^Department of Computer Science and Engineering, Korea University, Seoul, Korea. ^43^Center for Advanced Computation, Korea Institute for Advanced Study, Seoul, Korea. ^44^Interdisciplinary Graduate Program in Bioinformatics, Korea University, Seoul, Korea. ^45^Community High School, 401 N Division St, Ann Arbor, MI, 48104. ^46^National Center for Biotechnology Information, National Institute of Health (NCBI/NLM/NIH), USA. ^47^Biomolecular Interactions, Max Planck Institute for Developmental Biology, Spemannstr. 38, 72076 Tuebingen, Germany. ^48^Quantitative Biology Center, University of Tuebingen, Auf der Morgenstelle 8, 72076 Tuebingen, Germany. ^49^Data Science Program, Worcester Polytechnic Institute, Worcester, MA, USA. ^50^Department of Computer Science, Worcester Polytechnic Institute, Worcester, MA, USA. ^51^Department of Medicine, College of Physicians & Surgeons, Columbia University, New York, NY, USA. ^52^Qatar Computing Research Institute, Hamad Bin Khalifa University, Doha, Qatar. ^53^Institute of Social and Preventive Medicine (IUMSP), Lausanne University Hospital, Lausanne, Switzerland. ^54^Department of Surgery, Massachusetts General Hospital, Harvard Medical School, Boston, Massachusetts, USA. ^55^Institute for Biological Psychiatry, Mental Health Center Sct. Hans, University of Copenhagen, Roskilde, Denmark. ^56^Stanley Center at the Broad Institute of MIT and Harvard, Cambridge, Massachusetts, USA. ^57^Verge Genomics, San Francisco, CA, USA. ^58^Division of Geriatrics, Department of Medicine, University of California, San Francisco, USA. ^59^Centre for Cancer Biology, University of South Australia. ^60^School of Information Technology and Mathematical Sciences, University of South Australia. ^61^Department of Chemistry, Kangwon National University, 1 Kangwondaehak-gil, Chuncheon, 24341, Republic of Korea. ^62^BlueSkyIt, Amsterdam, the Netherlands. ^63^The Liver Care Center and Divisions of Gastroenterology, Hepatology and Nutrition, Cincinnati Children’s Hospital Medical Center, Cincinnati, OH, USA. ^64^Department of Computer Science and Engineering, University at Buffalo, Buffalo, NY, USA. ^65^Dept. of Env. Health, Division of Biostatistics and Bioinformatics, University of Cincinnati, OH, USA. ^66^Broad Institute of Harvard and MIT, Cambridge, MA. ^67^Bill and Melinda Gates Foundation. ^68^School of Electrical Engineering and Computer Science, Queensland University of Technology, Brisbane, Australia. ^69^Dept. of Electrical & Computer Engineering, Texas A&M University, USA. ^70^Department of Computer Science and Engineering, Indian Institute of Technology Madras, Chennai, India. ^71^Rockville, MD, USA (No affiliation). ^72^European Molecular Biology Laboratory, European Bioinformatics Institute (EMBL-EBI), Wellcome Genome Campus, Cambridge CB10 1SD, UK. ^73^RWTH Aachen University, Faculty of Medicine, Joint Research Centre for Computational Biomedicine, 52057 Aachen, Germany. ^74^Blavatnik School of Computer Science, Tel Aviv University, Tel Aviv 69978, Israel. ^75^Department of Biostatistics, the University of Texas MD Anderson Cancer Center, Houston, TX, USA. ^76^Sage Bionetworks, Seattle, Washington 98109, USA. ^77^Institute of Molecular Life Sciences and Swiss Institute of Bioinformatics, University of Zurich, Zurich, Switzerland. ^78^Memphis, TN, USA (No affiliation). ^79^CREST, JST, Tokyo, Japan. ^80^Department of Medical Science Mathematics, Medical Research Institute, Tokyo Medical and Dental University, Tokyo, Japan. ^82^ProGeLife, Marseille, France. ^83^Disease Science & Technology, Biocon Bristol-Myers Squibb Research Centre, Bangalore, India. ^84^Broad Institute of Harvard and MIT, Cambridge, MA. ^85^Janssen Research and Development. ^86^Department of Psychiatry, Yale School of Medicine, West Haven, CT, USA. ^87^Institute of Intelligent Machines, Hefei Institutes of Physical Science, Chinese Academy of Sciences, Hefei, Anhui, China. ^88^Department of Statistics and Operations Research, University of North Carolina at Chapel Hill, Chapel Hill, NC, USA. ^89^Key Laboratory of Systems Biology, Institute of Biochemistry and Cell Biology, Shanghai Institutes for Biological Sciences, Chinese Academy of Sciences. ^90^Computational biology consulting, avenue Kleber 100, Paris, France. ^91^School of Engineering, Dali University. ^92^Department of Electrical Engineering, Stanford University, USA. ^93^Institute for Immunology, Perelman School of Medicine at the University of Pennsylvania, Philadelphia, Pennsylvania, USA. ^94^Epigenetics Institute, Perelman School of Medicine at the University of Pennsylvania, Philadelphia, Pennsylvania, USA. ^95^Department of Mathematics and Statistics, The University of the West Indies, Saint Augustine, Trinidad and Tobago. ^96^Roche Pharma Research and Early Development, Pharmaceutical Sciences, Roche Innovation Center Basel, F. Hoffmann-La Roche Ltd, 4070 Basel, Switzerland.

## Author contributions

S.C., D.L., Z.K., G.S., J.M., K.L., J.S.-R., S.B. and D.M. conceived the challenge; S.C., G.S., J.S.-R., S.B. and D.M. organized the challenge; S.C. and D.M. performed team scoring; S.C., M.E.A., J.C., M.T., T.F., J.D.Z., D.K.S., L.J.C. and D.M. analyzed results; J.M., T.N., R.N., A.S., K.L. and J.S.-R. constructed networks; J.C., J.L., B.H., X.H., D.K.S. and L.J.C. designed the top-performing method; the DREAM Module Identification Consortium provided data and performed module identification; S.B. and D.M. designed the study; and D.M. prepared the manuscript. All authors discussed the results and implications, and commented on the manuscript at all stages.

## Acknowledgments

The challenge was hosted on Sage Bionetwork’s Synapse platform (https://synapse.org/). The computations were performed at the Vital-IT (http://www.vital-it.ch) Center for high-performance computing of the SIB Swiss Institute of Bioinformatics. This work was supported by the Swiss National Science Foundation (grant FN 310030_152724/1 to S.B. and grant FN 31003A-169929 to Z.K.), SystemsX.ch (grant SysGenetiX to S.B. and grant AgingX to Z.K.), the Swiss Institute of Bioinformatics (Z.K. and S.B.), the Leenaards Foundation (Z.K.), the US National Science Foundation (grant DMS-1812503 to L.C. and X.H.), and the National Institutes of Health (grant R01 HD076140 to D.K.S.).

## Methods

### Network compendium

A collection of six gene and protein networks for human were provided by different groups for this challenge. The two protein-protein interaction and signaling networks are custom or new versions of existing interaction databases that were not publicly available at the time of the challenge. The remaining networks were yet unpublished at the time of the challenge. This was important to prevent participants from deanonymizing challenge networks by aligning them to the original networks. The original networks, anonymized networks and the mappings from gene symbols to anonymized IDs are available on the challenge website.

Networks were released for the challenge in anonymized form. Anonymization consisted in replacing the gene symbols with randomly assigned ID numbers. In Sub-challenge 1 each network was anonymized individually, i.e., node *k* of network A and node *k* of network B are generally not the same genes. In Sub-challenge 2 all networks were anonymized using the same mapping, i.e., node *k* of network A and node *k* of network B are the same gene. Since the networks were unpublished, it was practically impossible for participants to infer the gene identities. Participants also agreed not to attempt to infer gene identities as part of the challenge rules.

All networks are undirected and weighted, except for the signaling network, which is directed and weighted. Basic properties and similarity between the networks are shown in Figs. 1A and **S2E**. Below we briefly summarize each of the six networks. Detailed descriptions of networks 4, 5 and 6 are available on GeNets (Li et al., 2018), a web platform for network-based analysis of genetic data (http://apps.broadinstitute.org/genets).

#### Network 1: STRING protein-protein interaction network

The first network was obtained from STRING, a database of known and predicted protein-protein interactions (Szklarczyk et al., 2015). STRING includes aggregated interactions from primary databases as well as computationally predicted associations. Both physical protein interactions (direct) and functional associations (indirect) are included. The challenge network corresponds to the human protein-protein interactions of STRING version 10.0, where interactions derived from text-mining were removed. Edge weights correspond to the STRING association score after removing evidence from text mining. The network was provided by Damian Szklarczyk and Christian von Mering (University of Zürich).

#### Network 2: InWeb protein-protein interaction network

The second network is the InWeb protein-protein interaction network (Li et al., 2017). InWeb aggregates physical protein-protein interactions from primary databases and the literature. The challenge network corresponds to InWeb version 3. Edge weights correspond to a confidence score that integrates the evidence of the interaction from different sources.

#### Network 3: OmniPath signaling network

The third network is the OmniPath signaling network (Türei et al., 2016). OmniPath integrates literature-curated human signaling pathways from 27 different sources, of which 20 provide causal interaction, 7 deliver undirected interactions. These data were integrated to form a directed weighted network. The edge weights correspond to a confidence score that summarizes the strength of evidence from the different sources.

#### Network 4: GEO co-expression network

The fourth network is a co-expression network based on Affymetrix HG-U133 Plus 2 arrays extracted from the Gene Expression Omnibus (GEO) (Barrett et al., 2011). In order to adjust for non-biological variation, data were rescaled by fitting a loess-smoothed power law curve to a collection of 80 reference genes (ten sets of ~8 genes each, representing different strata of expression) using nonlinear least squares regression within each sample. All samples were then quantile normalized together as a cohort. This approach is described fully in (Subramanian et al., 2017). After filtering out samples that did not pass quality control, a gene expression matrix of 22,268 probesets by 19,019 samples was obtained. Probes were mapped to genes by averaging and the pairwise Spearman correlation of genes across samples was computed. The matrix was thresholded to include the top 1M strongest positive correlations resulting in an undirected, weighted network. The edge weights correspond to the correlation coefficients.

#### Network 5: Achilles cancer co-dependency network

The fifth network is a functional gene network derived from the Project Achilles dataset v2.4.3 (Cowley et al., 2014). Project Achilles performed genome-scale loss-of-function screens in 216 cancer cell lines using massively parallel pooled shRNA screens. Cell lines were transduced with a library of 54,000 shRNAs, each targeting one of 11,000 genes for RNAi knockdown (~5 shRNAs per gene). The proliferation effect of each shRNA in a given cell line could be assessed using Next Generation Sequencing. From these data, the dependency of a cell line on each gene (the gene essentiality) was estimated using the ATARiS method. This led to a gene essentiality matrix of 11,000 genes by 216 cell lines. Pairwise correlations between genes were computed and the resulting co-dependency network was thresholded to the top 1M strongest positive correlations, analogous to how the co-expression network was constructed. Project Achilles data was kindly provided by Aviad Tsherniak and Barbara Weir (Broad Institute).

#### Network 6: CLIME homology-based network

The sixth network is a functional gene network based on phylogenetic relationships identified using the CLIME (clustering by inferred models of evolution) algorithm (Li et al., 2014). CLIME can be used to expand pathways (gene sets) with additional genes using an evolutionary model. Briefly, given a eukaryotic species tree and homology matrix, the input gene set is partitioned into evolutionarily conserved modules (ECMs), which are then expanded with new genes sharing the same evolutionary history. To this end, each gene is assigned a log-likelihood ratio (LLR) score based on the ECMs inferred model of evolution. CLIME was applied to 1,025 curated human gene sets from GO and KEGG using a 138 eukaryotic species tree, which resulted in 13,307 expanded ECMs. The network was constructed by adding an edge between every pair of genes that co-occurred in at least one ECM. Edge weights correspond to the mean LLR scores of the two genes.

### Challenge structure

Participants were challenged to apply network module identification methods to predict functional modules (gene sets) based on network topology. Valid modules had to be non-overlapping (a given gene could be part of either zero or one module, but not multiple modules) and comprise between 3 and 100 genes. Modules did not have to cover all genes in a network. The number of modules per network was not fixed: teams could submit any number of modules for a given network (the maximum number was limited due to the fact that modules had to be non-overlapping). In Sub-challenge 1, teams were required to submit a separate set of modules for each of the six networks. In Sub-challenge 2, teams were required to submit a single set of modules by integrating information across multiple networks (it was permitted to use only a subset of the six networks).

The challenge consisted of a leaderboard phase and the final evaluation. The leaderboard phase was organized in four rounds, where teams could make repeated submissions and see their score on each network. Due to the high computational cost of scoring the module predictions on a large number of GWAS datasets (see next section), a limit for the number of submissions per team was set in each round taking into consideration our computational resources and the number of participating teams. The total number of submissions that any given team could make over the four leaderboard rounds was thus limited to only 25 and 41 for the two sub-challenges, respectively. For the final evaluation, a single submission including method descriptions and code was required per team, which was scored on a separate set of GWASs after the challenge closed to determine the top performers.

The submission format and rules are described in detail on the challenge website (https://www.synapse.org/modulechallenge).

### Challenge scoring

We have developed a novel framework to empirically assess module identification methods on molecular networks using GWAS data. In contrast to functional gene annotations and pathway databases such as GO, which sometimes originate from similar types of functional genomics data as the network modules, GWAS data are orthogonal to the networks and thus provide an independent means of validation. In order to cover diverse molecular processes, we compiled a large collection of 180 GWAS datasets from public sources. The collection was split into two sets of 76 and 104 GWASs used for the leaderboard phase and the final evaluation, respectively (**Table S1**).

#### Gene and module scoring using Pascal

SNP-trait association p-values from a given GWAS were integrated across genes and modules using the Pascal (pathway scoring algorithm) tool (Lamparter et al., 2016). Briefly, Pascal combines analytical and numerical solutions to efficiently compute gene and module scores from SNP p-values, while properly correcting for linkage disequilibrium (LD) correlation structure prevalent in GWAS data. To this end, LD information from a reference population is used (here, the European population of the 1000 Genomes Project was employed as we only included GWASs with predominantly European cohorts). Compared to alternative gene scoring methods that rely on Monte Carlo simulations, Pascal is about 100 times faster and more precise (Lamparter et al., 2016). The fast gene scoring is critical as it allows module genes that are in LD, and can thus not be treated independently, to be dynamically rescored. This amounts to fusing the genes of a given module that are in LD and computing a new score that takes the full LD structure of the corresponding locus into account. Finally, Pascal tests modules for enrichment in high-scoring (potentially fused) genes using a modified Fisher method, which avoids any p-value cutoffs inherent to standard binary enrichment tests. As background gene set, the genes of the given network were used. Lastly, the resulting nominal module p-values were adjusted to control the FDR via the Benjamini-Hochberg procedure. A snapshot of the Pascal version used for the challenge is available on the challenge website.

#### Scoring metric

In Sub-challenge 1, the score for a given network was defined as the number of modules with significant Pascal p-values at a given FDR cutoff in at least one GWAS (called trait-associated modules). Thus, modules that were hits for multiple GWAS traits were only counted once. The overall score was defined as the sum of the scores obtained on the six networks (i.e., the total number of trait-associated modules across all networks). For the official challenge ranking a 5% FDR cutoff was defined, but performance was further reported at 10%, 2.5% and 1% FDR.

Prior to the challenge, we performed an analysis to explore whether this scoring metric would favor a particular resolution for modules and thus bias results, e.g., towards decomposing modules into a larger number of small sub-modules. To this end, we applied off-the-shelf community detection methods and varied parameters to obtain different number and size of modules. We found no systematic bias in the scores for a specific module granularity and this result was confirmed in the challenge (see Section “Complementarity of different module identification approaches and Fig. S3B-E). The key element of the scoring function that was designed to fairly assess module collections with different average module sizes was the higher multiple testing burden applied when a larger number of smaller modules was submitted.

Module predictions in Sub-challenge 2 were scored using the exact same methodology and FDR cutoffs. The only difference to Sub-challenge 1 was that submissions consisted of a single set of modules (instead of one for each network) and there was thus no need to define an overall score. As background gene set, the union of all genes across the six networks was used.

#### Robustness analysis of challenge ranking

To gain a sense of the robustness of the ranking with respect to the GWAS data, we subsampled the set of 104 GWASs used for the final evaluation (called the “test set”) by drawing 76 GWASs (same number of GWASs as in the leaderboard set; note that we have to do subsampling rather than resampling of GWASs because the scoring counts the number of modules that are associated to at least one GWAS, i.e., including the same GWASs multiple times does not affect the score). We applied this approach to create 1,000 subsamples of the test set. The methods were then scored on each subsample.

The performance of every method *m* was compared to the highest-scoring method across the subsamples by the paired Bayes factor *K_m_*. That is, the method with the highest overall score in the test set (all 104 GWASs) was defined as reference (i.e., method *K1* in Sub-challenge 1). The score *S(m, k)* of method *m* in subsample *k* was thus compared with the score *S(ref, k)* of the reference method in the same subsample *k*. The Bayes factor *K_m_* is defined as the number of times the reference method outperforms method *m*, divided by the number of times method *m* outperforms or ties the reference method over all subsamples. Methods with *K_m_* < 3 were considered a tie with the reference method (i.e., method *m* outperforms the reference in more than 1 out of 4 subsamples).

### Module identification methods

Here we provide an overview of module identification approaches applied in the two sub-challenges, including a detailed description of the top-performing method. Full descriptions and code of all methods are available on the challenge website (https://www.synapse.org/modulechallenge).

#### Overview of module identification methods in Sub-challenge 1

Based on descriptions provided by participants, module identification methods were classified into different categories (Fig. 2A). Categories and corresponding module identification methods are summarized in Table 1. In the following, we first give an overview of the different categories and top-performing methods, and then describe common pre- and post-processing steps used by these methods:

- **Kernel clustering**. Instead of working directly on the networks themselves, these methods cluster a kernel matrix, where each entry (*i*, *j*) of that matrix represents the closeness of nodes *i* and *j* in the network according to the particular similarity function, or kernel that was applied. Some of the kernels that were applied are well-known for community detection, such as the exponential diffusion kernel based on the graph Laplacian (Kondor and Lafferty, 2002) employed by method *K6*. Others, such as the LINE embedding algorithm (Tang et al., 2015) employed by method *K3* and the kernel based on the inverse of the weighted diffusion state distance (Cao et al., 2013, 2014) employed by method *K1*, were more novel. Method *K1* was the best-performing method of the challenge and is described in detail below.
- **Modularity optimization.** This method category was, along with random-walk-based methods (see below), the most popular type of method contributed by the community. Modularity optimization methods use search algorithms to find a partition of the network that maximizes the modularity *Q* (commonly defined as the fraction of within-module edges minus the expected fraction of such edges in a random network with the same node degrees) (Newman and Girvan, 2004). The most popular algorithm was Louvain community detection (Blondel et al., 2008). At least eight teams employed this algorithm in some form as either their main method or one of several methods. The top team of the category (method *M1*), which ranked second overall, first sparsified networks by removing low confidence edges. A mixture of several established community detection algorithms was then employed in order to search for a partition that optimized modularity. Importantly, these algorithms were extended with an additional resistance parameter that penalized merging of communities (Arenas et al., 2008); increasing the resistance parameter thus led to partitions with a larger number of communities. Communities above the size limit (100 nodes) were subdivided recursively by reapplying the same community detection algorithms to the corresponding subnetworks (see below).
- **Random-walk-based methods.** These methods take inspiration from random walks or diffusion processes over the network. Several teams used the established Walktrap (Pons and Latapy, 2005) and Infomap (Rosvall et al., 2009) algorithms. The top team of this category (method *R1*) used a sophisticated random-walk method based on multi-level Markov clustering (Satuluri et al., 2010). The method modifies basic Markov Clustering in two ways. First, a hierarchical view of the graph is considered by successively coarsening neighborhoods into fewer supernodes. The clustering is first run on the coarsened graph, enabling the detection of communities at varying scales. Second, a balance parameter is introduced that adjusts for nodes to preferentially join smaller communities, thus leading to more balanced community sizes. Similar to method *M1* described above, networks were first sparsified and communities above the size limit were recursively subdivided. While we did not include kernel methods in the “random walk” category, several of the successful kernel clustering methods used random-walk-based measures within their kernel functions.
- **Local methods**. Only three teams used local community detection methods, including agglomerative clustering and seed set expansion approaches. The top team of this category (method *L1*) first converted the adjacency matrix into a topology overlap matrix (Ravasz et al., 2002), which measures the similarity of nodes by their topological overlap based on the number of neighbor they have in common. The team then used the SPICi algorithm (Jiang and Singh, 2010), which iteratively adds adjacent genes to cluster seeds such as to improve their local density.
- **Hybrid methods**. Seven teams employed hybrid methods that leveraged clusterings produced by several of the different main approaches listed above. These teams applied more than one community detection method to each network in order to get larger and more diverse sets of predicted modules. The most common methods applied were Louvain (Blondel et al., 2008) hierarchical clustering, and Infomap (Rosvall et al., 2009). Two different strategies were used to select a final set of modules for submission: (1) choose a single method for each network according to performance in the leaderboard round, and (2) select modules from all applied methods according to a topological quality score such as the modularity or conductance (Fortunato and Hric, 2016).
- **Ensemble methods.** Much like hybrid methods, ensemble methods leverage clusterings obtained from multiple community detection methods (or multiple stochastic runs of a single method). However, instead of selecting individual modules according to a quality score, ensemble methods merge alternative clusterings to obtain potentially more robust consensus predictions (Lancichinetti and Fortunato, 2012). Our method to derive consensus module predictions from team submissions is an example of an ensemble approach (described in detail below).

Besides the choice of the community detection algorithm, there are other steps that critically affected performance, including pre-processing of the network data, setting of method parameters, and post-processing of predicted modules. We describe successful approaches employed by challenge participants to address these issues below (pre- and post-processing steps of challenge methods are also summarized in Table 1):

- **Pre-processing**. Data pre-processing often plays a key role in the analysis of noisy data, such as biological network data. Most networks in the challenge were densely connected, including many edges of low weight that are likely noisy. Some of the top teams (e.g., *M1*, *R1*, *L1*) benefitted from sparsifying these networks by discarding weak edges before applying their community detection methods. An added benefit of sparsification is that it typically reduces computation time. Few teams also normalized the edge weights of a given network to make them either normally distributed or fall in the range between zero and one. Not all methods required pre-processing of networks, for example the top performing method (*K1*) was applied to the original networks without any sparsification or normalization steps.
- **Parameter setting**. Most community detection methods have parameters that need to be specified, typically to control the resolution of the clustering (the number and size of modules). While some methods have parameters that explicitly set the number of modules (e.g., the top-performing method *K1*), other methods have parameters that indirectly control the resolution (e.g., the resistance parameter of the runner-up method *M1*). Teams used the leaderboard phase to optimize the parameters of their method. Note that teams could make at most 25 submissions during the leaderboard phase, which limited the parameter space that could be explored in particular for methods with multiple parameters. While there were also methods that had no parameters to set (e.g., the classic Louvain algorithm), these methods have an intrinsic resolution that may not always be optimal for a given network and target application.
- **Post-processing**. Depending on the target application, the output of community detection methods may need to be post-processed. In biological networks, most methods typically lead to highly imbalanced module sizes. That is, some modules may be very small (e.g., just one or two genes), while others are extremely large (e.g., thousands of genes). Both extremes are generally not useful to gain biological insights at the pathway level. In the challenge, module sizes were thus required to be between 3 and 100 genes. Since current community detection methods generally do not allow such constraints on module size to be specified, teams used different post-processing steps to deal with modules outside of this range. A successful strategy employed by teams to break down large modules was to recursively apply their method to each of these modules. Alternatively, all modules of invalid size were merged and the community detection method was re-applied to the corresponding subnetwork. Finally, modules with less than three genes were often discarded (i.e., the corresponding genes were not included in any of the submitted modules). Some teams also discarded larger modules that were deemed low quality according to a topological metric, although this strategy was generally not beneficial.

#### Top-performing team method

The top-performing team developed a kernel clustering approach (method *K1*) based on a distance measure called Diffusion State Distance (DSD) (Cao et al., 2013, 2014), which they further improved for this challenge (Crawford et al., in preparation). DSD produces a more informative notion of proximity than the typical shortest path metric, which measures distance between pairs of nodes by the number of hops on the shortest path that joins them in the network. More formally, consider the undirected network *G*(*V*, *E*) on the node set *V* = {*v*_1_,*v*_2_,*v*_3_,…,*v*_*n*_} with |*V*|= *n*. *He*^*t*^(*v*_*x*_, *v*_*y*_) is defined as the defined as the expected number of times that a random walk (visiting neighboring nodes in proportion to their edge weights) starting at node *v*_*x*_ and proceeding for some fixed t steps will visit node *v*_*y*_ (the walk includes the starting point, i.e., 0th step). Taking a global view, we define the n-dimensional vector *He*^*t*^(*v*_*x*_) whose *i*th entry is the *He*^*t*^(*v*_*x*_, *v*_*i*_) value to network node *v*_*i*_. Then the *DSD*^*t*^ distance between two nodes *v*_*x*_ and *v*_*y*_ is defined as the *L*1 norm of the difference of their *He*^*t*^ vectors, i.e.

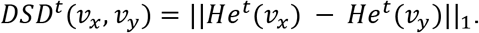

It can be shown that DSD is a metric and converges as *t* → ∞, allowing DSD to be defined independently from the value *t* (Cao et al., 2013). The converged DSD matrix can be computed tractably, with an eigenvalue computation, as

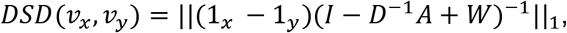

where *D* is the diagonal degree matrix, *A* is the adjacency matrix, and *W* is the matrix where each row is a copy of *π*, the degrees of each of the nodes, normalized by the sum of all the vertex degrees (in the unweighted case; weighted edges can be normalized proportional to their weight), and 1_x_ and 1_y_ are the vectors that are zero everywhere except at position x and y, respectively. The converged DSD matrix was approximated using algebraic multigrid techniques (Crawford et al., in preparation). Note that for the signaling network, edge directions were kept and low-weight back edges were added so that the network was strongly connected; i.e. if there was a directed edge from *v*_*x*_ to *v*_*y*_, an edge from *v*_*y*_ to *v*_*x*_ of weight equal to 1/100 of the lowest edge weight in the network was added.

A spectral clustering algorithm (Ng et al., 2001) was used to cluster the DSD matrix of a given network. Note that the spectral clustering algorithm operates on a similarity matrix (i.e., entries that are most alike have higher values in the matrix). However, the DSD matrix is a distance matrix (i.e., similar entries have low DSD values). The radial basis function kernel presents a standard way to convert the DSD matrix to a similarity matrix; it maps low distances to high similarity scores and vice-versa. Since the spectral clustering algorithm employed uses k-means as the underlying clustering mechanism, it takes a parameter k specifying the number of cluster centers. The leaderboard rounds were utilized to measure the performance of different k. Also note that spectral clustering produces clusters of size less than 3, and clusters of size more than 100. Whenever a cluster of size less than 3 was produced, those vertices were not included in any cluster for that network. Whenever a cluster of size more than 100 was produced, spectral clustering was called recursively to split that cluster into two subclusters (i.e., k=2) until all clusters were of size < 100.

The top-performing team also used a different algorithm to search for dense bipartite subgraph module structure in half of the challenge networks and merged these modules (which were rare) with the clusters generated by their main method (Gallant et al., 2013). However, a post-facto analysis of their results showed that this step contributed few modules and the score would have been similar with this additional procedure omitted (see method description on Challenge website for details).

#### Overview of module identification methods in Sub-challenge 2

In Sub-challenge 2, few teams employed dedicated multi-network community detection methods (De Domenico et al., 2015; Didier et al., 2015). The majority of teams first built an integrated network by merging either all six or a subset of the challenge networks, and then applied single-network methods (typically the same method as in Sub-challenge 1) to modularize the integrated network. For example, the team with highest score in Sub-challenge 2 merged the two protein interaction networks and then applied the Louvain algorithm to identify modules in the integrated network. The top performing team from Sub-challenge 1 also performed competitively in Sub-challenge 2. They applied their single-network method (*K1*) to an integrated network consisting of the union of all edges from the two protein interaction networks and the coexpression network.

Similar to Sub-challenge 1, teams used the leaderboard phase to set parameters of their methods. However, besides the parameters of the community detection method, there were additional choices to be made, whether to use all or only a subset of the six networks and how to integrate them.

### Consensus module predictions

We developed an ensemble approach to derive consensus modules from a given set of team submissions (see Fig. S2A for a schematic overview). In Sub-challenge 1, a consensus matrix *C^n^* was defined for each network *n*, where each element *c_ij_* corresponds to the fraction of teams that put gene *i* and *j* together in the same module in this network. That is, *c_ij_* equals one if all teams clustered gene *i* and *j* together, and *cij* equals zero if none of the teams clustered the two genes together. The top-performing module identification method (*K1*) was used to cluster the consensus matrix (i.e., the consensus matrix was considered a weighted adjacency matrix defining a functional gene network, which was clustered using the top module identification method of the challenge). Method *K1* has only one parameter to set, which is the number of cluster centers used by the spectral clustering algorithm (see previous section). This parameter was set to the median number of modules submitted by the considered teams for the given network. The consensus module predictions described in the main text were derived from the submissions of the top 50% teams (i.e., 21 teams) with the highest overall score on the leaderboard GWAS set. (Results for different cutoffs regarding the percentage of teams included are reported in Fig. S2C.)

Multi-network consensus modules were obtained by integrating team submissions from Sub-challenge 1 across all six networks using the same approach (see Fig. S2B). The same set of teams was considered (i.e., top 50% on the leaderboard GWAS set). First, a multi-network consensus matrix was obtained by taking the mean of the six network-specific consensus matrices *C^n^*. The multi-network consensus matrix was then clustered using method *K1* as described above, where the number of cluster centers was set to the median number of modules submitted by the considered teams across all networks.

Two additional, more sophisticated approaches to construct consensus matrices *C^n^* were tested: (1) normalization of the contribution of each module by the module size led to similar results as the basic approach described above, and (2) unsupervised estimation of module prediction accuracy using the Spectral Meta Learner ensemble method (Parisi et al., 2014) did not perform well in this context (Fig. S2D).

### Similarity of module predictions

To define a similarity metric between module predictions from different methods, we represented module predictions as vectors. Namely, the set of modules predicted by method *m* in network *k* was represented as a prediction vector of *P*_*mk*_ length *N*_*k*_(*N*_*k*_ − 1)/2, where *N*_*k*_ is the number of genes in the network. Each element of this vector corresponds to a pair of genes and equals 1 if the two genes are in the same module and 0 otherwise. Accordingly, for any two module predictions (method *m*_1_ applied to network *k*_1_, and method *m*_2_ applied to network *k*_2_), we calculated the distance as follows:

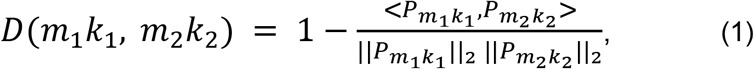

where <.,.> is the Euclidean inner product, ‖.‖_2_ is the Euclidean norm, and *D* is the (symmetric) distance matrix between the 252 module predictions submitted in Sub-challenge 1 (i.e., 42 methods applied to each of six networks). The distance matrix *D*was used as input to the Multidimensional Scaling (MDS) analysis for dimensionality reduction in Fig. 3A.

Similarity between method predictions across networks was calculated in the same way. To this end, the prediction vectors *P*_*mk*_ of method *m* for the six networks (*k* = 1,2,…,6) were concatenated, forming a single vector *P*_*m*_ that represents the module predictions of that method for all six networks. A corresponding distance matrix between the 42 methods was computed using the same approach as described above (Equation 1) and used as input for hierarchical clustering in Fig. S3A.

### Overlap between trait-associated modules

Three different metrics were considered to quantify the overlap between trait-associated modules from different methods and networks. The first metric was the Jaccard index, which is defined as the size of the intersection divided by the size of the union of two modules (gene sets) *A* and *B*:

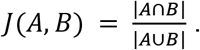

The Jaccard index measures how similar two modules are, but does allow the detection of sub-modules. For example, consider a module *A* of size 10 that is a submodule of a module *B* of size 100. In this case, even though 100% of genes of the first module are comprised in the second module, the Jaccard index is rather low (0.1). To capture sub-modules, we thus considered in addition the percentage of genes of the first module that are comprised in the second module:

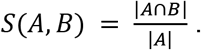

Lastly, we also evaluated the significance of the overlap. To this end, we computed the p-value *p*_*AB*_ for the overlap between the two modules using the hypergeometric distribution. P-values were adjusted using Bonferroni correction given the number of module pairs tested.

Based on these three metrics, we categorized the type of overlap that a given trait-module *A* had with another trait-module *B* as:

1. *strong overlap* if *J*(*A*, *B*) ≥ 0.5 and *p*_*AB*_ < 0.05;
2. *submodule* if *J*(*A*, *B*) < 0.5 and *S*(*A*, *B*) − *J*(*A*, *B*) ≥ 0.5 and *p*_*AB*_ < 0.05;
3. *partial overlap* if *J*(*A*, *B*) < 0.5 and *S*(*A*, *B*) − *J*(*A*, *B*) < 0.5 and *p*_*AB*_ < 0.05;
4. *insignificant overlap* if *p*_*AB*_ ≥ 0.05.

This categorization was used to get a sense of the type of overlap between trait modules from all methods (see Fig. 3B).

### Trait similarity network

We defined a network level similarity between GWAS traits based on overlap between trait-associated modules. To this end, we only considered the most relevant networks for our collection of GWAS traits, i.e., the two protein interaction, the signaling and the co-expression network (see Fig. 2D). For a given network, the set of “gtrait-module genes” *G*_*T*_ was obtained for every trait *T*by taking the union of the modules associated with that trait across all challenge methods. (If different GWASs were available for the same trait type (see **Table S1**), the union of all corresponding trait-associated modules was taken). The overlap between every pair of trait-module gene sets 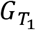 and 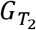 was evaluated using the Jaccard index 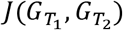 and the hypergeometric p-value 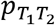 as described in the previous section. P-values were adjusted using Bonferroni correction. For the visualization as a trait-trait network in Fig. 4C, an edge between traits *T*_1_ and *T*_2_ was added if the overlap was significant 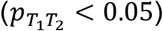 in at least three out of the four considered networks, and node sizes and edge weights were set proportional to the average number of trait-module genes and the average Jaccard index across the four networks, respectively.

### Evaluation of candidate trait genes

Trait-associated modules comprise many genes that show only borderline or no signal in the corresponding GWAS (called “candidate trait genes”). To assess whether modules correctly prioritized candidate trait genes, we considered eight traits for which older (lower-powered) and more recent (higher-powered) GWAS datasets were available in our test set (Fig. S4A). This allowed us to evaluate how well trait-associated modules and candidate trait genes predicted using the lower-powered GWAS datasets were supported in the higher-powered GWAS datasets.

We only considered candidate trait genes that were predicted solely because of their membership in a trait-associated module, i.e., that did not show any signal in the lower-powered GWAS as defined by: (i) a high gene p-value (*p* > 1E-4, i.e., two orders of magnitude above the genome-wide significance threshold of 1E-6) and (ii) genomic location of more than one megabase away from the nearest significant locus of the corresponding GWAS. Gene p-values were computed using Pascal as described above (see “Gene and module scoring using the Pascal tool”). Finally, the Pascal p-value of all candidate trait genes was evaluated for the higher-powered GWAS. Since there is a genome-wide tendency for p-values to become more significant in higher-powered GWAS data (Boyle et al., 2017), Pascal p-values were also evaluated for a background gene set (all genes that meet the two conditions (*i*, *ii*) but do not belong to trait-associated modules of the lower-powered GWAS). Fig. 5C shows the cumulative distribution of Pascal p-values for the candidate trait genes as well as the background genes.

### Functional enrichment analysis

In order to test network modules for enrichment in known gene functions and pathways, we considered diverse annotation and pathway databases. GO annotations for biological process, cellular component, and molecular functions were downloaded from the GO website (http://geneontology.org, accessed on January 20, 2017). Curated pathways (KEGG, Reactome, and BioCarta) were obtained from MSigDB version 5.2 (http://software.broadinstitute.org/gsea). We also created a collection of gene sets reflecting mouse mutant phenotypes, as defined by the Mammalian Phenotype Ontology (Blake et al., 2017). We started with data files HMD_HumanPhenotype.rpt and MGI_GenePheno.rpt, downloaded from the Mouse Genome Informatics database (http://www.informatics.jax.org) on February 21, 2016. The first file contains human-mouse orthology data and some phenotypic information; we then integrated more phenotypic data from the second file, removing the two normal phenotypes MP:0002169 (“no abnormal phenotype detected”) and MP:0002873 (“normal phenotype”). For each remaining phenotype, we then built a list of all genes having at least one mutant strain exhibiting that phenotype, which we considered as a functional gene set.

Annotations from curated databases are known to be biased towards certain classes of genes. For example, some genes have been much more heavily studied than others and thus tend to have more annotations assigned to them. This and other biases lead to an uneven distribution of the number of annotations per genes (annotation bias). On the other hand, the gene sets (modules) tested for enrichment in these databases typically also exhibit bias for certain classes of genes (selection bias) (Glass and Girvan, 2014; Young et al., 2010). Standard methods for GO enrichment analysis use the hypergeometric distribution (i.e., Fisher’s exact test), the underlying assumption being that, under the null hypothesis, each gene is equally likely to be included in the gene set (module). Due to selection bias, this is typically not the case in practice, leading to inflation of p-values (Glass and Girvan, 2014; Young et al., 2010). Following Young et al. (2010), we thus used the Wallenius non-central hypergeometric distribution to account for biased sampling. Corresponding enrichment p-values were computed for all network modules and annotation terms (pathways). The genes of the given network were used as a background gene set. For each network, module identification method, and annotation database, the *M* × *T* nominal p-values of the *M* modules and *T* annotation terms (pathways) were adjusted using Bonferroni correction.

## Data and software availability

Challenge data, results, and code are available from the challenge website (https://synapse.org/modulechallenge). This includes:

- Official challenge rules;
- Gene scores for the compendium of 180 GWASs used in the challenge plus 5 additional GWASs obtained after the challenge (GWAS SNP p-values are available upon request);
- The molecular network collection (anonymized and deanonymized versions);
- Module identification method descriptions and code provided by teams;
- The final module predictions of all teams for both sub-challenges;
- Consensus module predictions for both sub-challenges;
- Method scores at varying FDR cutoffs;
- Individual module scores for all GWASs;
- Enriched functional annotations for all modules (GO, mouse mutant phenotypes, and diverse pathway databases);
- A snapshot of the PASCAL tool and scoring scripts.

The latest version of PASCAL and the source code is also available from the PASCAL website (https://www2.unil.ch/cbg/index.php?title=Pascal) and GitHub (https://github.com/dlampart/Pascal).

## Supplementary Figures and Tables

**Figure S1:**
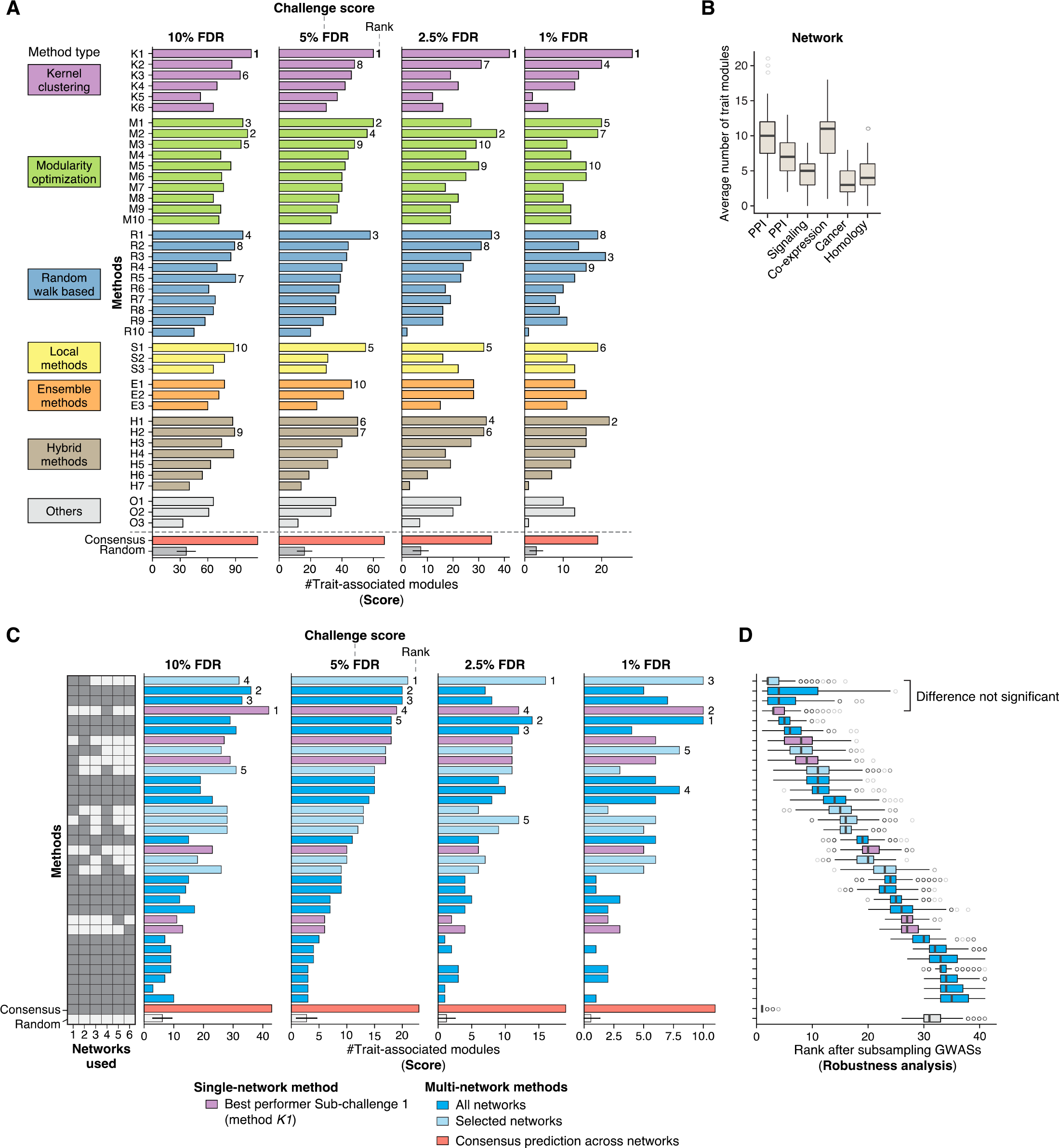
Assessment of Module Identification Methods. **(A)** Overall scores of the 42 module identification methods applied in Sub-challenge 1 at four different FDR cutoffs (10%, 5%, 2.5%, and 1% FDR). For explanation see legend of Fig. 2B, which shows the scores at 5% FDR (the predefined cutoff used for the challenge ranking). The top-performing method (*K1*) ranks first at all four cutoffs. The consensus prediction achieves the top score at 10% and 5% FDR, but not at the more stringent cutoffs. **(B)** Average number of trait-associated modules across all methods for each of the six networks. The most trait modules are found in the two protein-protein interaction (PPI) and the co-expression networks. Related to Fig. 2D, which shows the average number of trait modules relative to network size. **(C)** Final scores of multi-network module identification methods in Sub-challenge 2 at four different FDR cutoffs (10%, 5%, 2.5%, and 1% FDR). For explanation see legend of Fig. 3E, which shows the scores at 5% FDR (the predefined cutoff used for the challenge ranking). Ranks are indicated for the top five teams (ties are broken according to robustness analysis described in **Panel D**). The multi-network consensus prediction (red) achieves the top score at each FDR cutoff. Interestingly, the performance of methods integrating all five networks (dark blue) seems to drop substantially at the more stringent FDR thresholds. For example, the second and third ranking methods at both 5% and 10% FDR, which integrated all five networks, performed poorly at the 2.5% and 1% FDR thresholds (see second and third row from the top). This suggests that not only the absolute number of trait-associated modules, but also their quality in terms of association strength could not be improved by considering multiple networks. As mentioned in the Discussion, the challenge networks may not have been sufficiently related for multi-network methods to reveal meaningful modules spanning several networks. Indeed, the similarity between our networks in terms of edge overlap was small (Fig. S2E). Of note, there is an important conceptual difference between the multi-network methods that teams applied (blue) and the multi-network consensus prediction (red). While the former performed modularization on blended or multi-layer networks, the latter integrated the single-network module predictions obtained from each individual network (see Fig. S2B). Results thus suggest that our multi-network consensus approach is better suited than multi-layer module identification methods when network similarity is low. Exploring the performance of these different approaches when applied to networks of varying similarity is a promising avenue for future work. **(D)** Robustness of the overall ranking in Sub-challenge 2 was evaluated by subsampling the GWAS set used for evaluation 1,000 times. For each method, the resulting distribution of ranks is shown as a boxplot (using the 5% FDR cutoff for scoring). Related to Fig. 2C, which shows the same analysis for Sub-challenge 1. The difference between the top single-network module prediction and the top multi-network module predictions is not significant when sub-sampling the GWASs (Bayes factor < 3).

**Figure S2:**
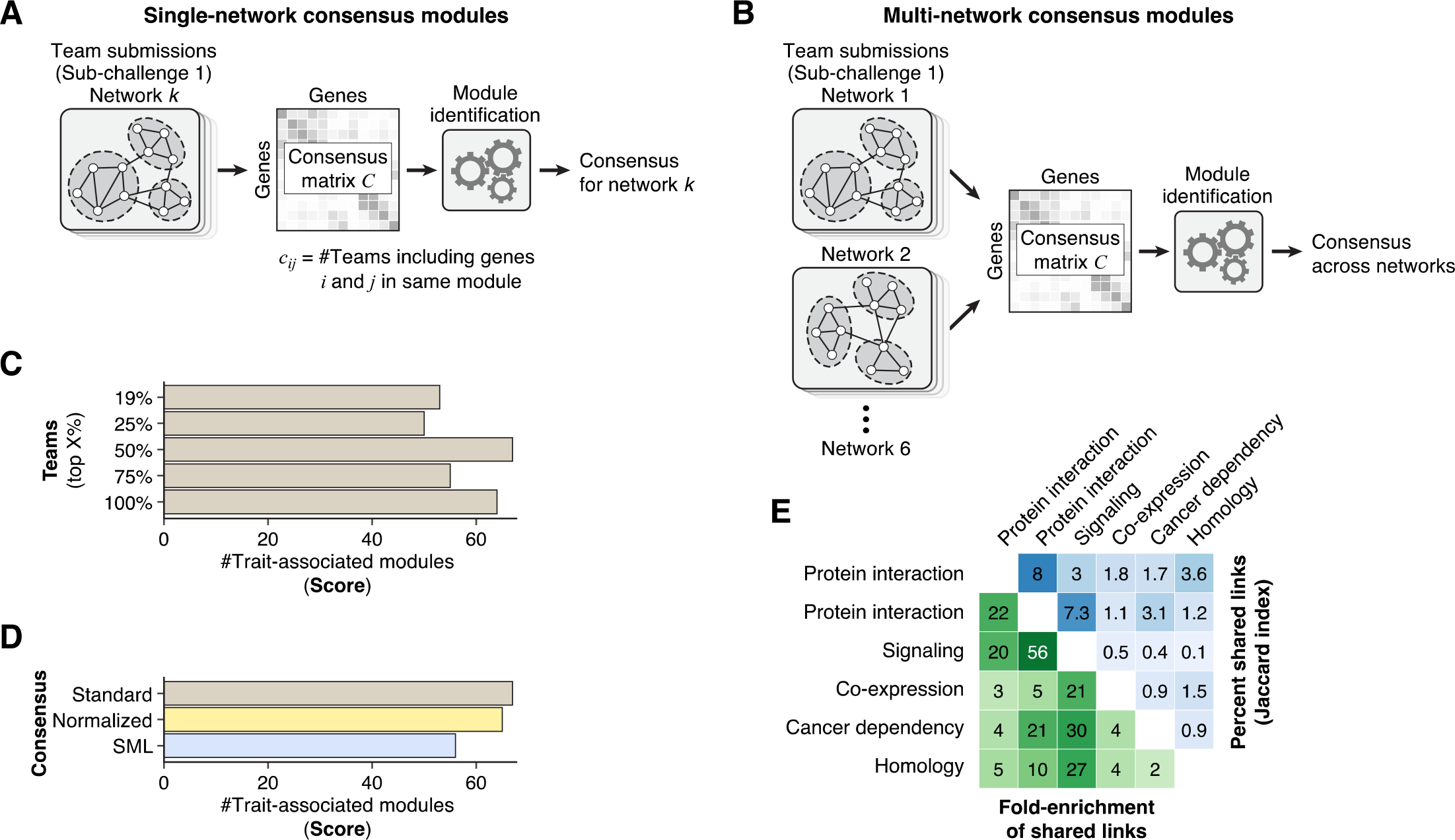
Consensus Module Predictions. **(A)** Schematic of the approach used to generate single-network consensus module predictions for Sub-challenge 1. For each network, module predictions from the top 50% of teams were integrated in a consensus matrix *C*, where each element *c_ij_* gives the fraction of teams that clustered gene *i* and *j* together in the same module in the given network (performance as the percentage of considered teams is varied is shown in **Panel C**). The overall score from the leaderboard round was used to select the top 50% of teams, i.e., the same set of teams was used for each network. The consensus matrix of each network was then clustered using the top-performing module identification method of the challenge (method *K1*; see Methods). **(B)** The approach used to generate multi-network consensus module predictions for Sub-challenge 2 was exactly the same as for single-network predictions, except that team submissions from all networks were integrated in the consensus matrix *C*. In other words, as input we still used the single-network predictions of the top 50% of teams from Sub-challenge 1, but instead of forming a consensus matrix for each network, a single cross-network consensus matrix was formed. This cross-network consensus matrix is then clustered using method *K1* as described above (see Methods). **(C)** Scores of the single-network consensus predictions as the percentage of integrated teams is varied. We considered the top 25%, 50%, 75% and 100% of teams, as well as the top eight (19%) teams (these are the teams that ranked 2nd, or tied with the team that ranked 2nd, at any of the considered FDR cutoffs). **(D)** Performance of different methods to construct the consensus matrix *C*. In addition to the basic approach described above (*Standard*), two more sophisticated approaches to construct the consensus matrix were evaluated (*Normalized* and *SML*). In each case, the same set of team submissions were integrated (top 50%) and method *K1* was applied to cluster the resulting consensus matrix. The first alternative (*Normalized*) is similar to the basic method but further assumes that appearing together in a smaller cluster is stronger evidence that a pair of genes is associated than appearing together in a larger cluster. Thus, each cluster’s contribution to the consensus matrix was normalized by the size of the cluster. Furthermore, we normalized the *ij*-entry of the consensus matrix by the number of methods that assigned gene *i* to a cluster, thus taking the presence of background genes into account. We found that the consensus still achieved the top score with these normalizations, but there was no improvement compared to the basic approach. The second method is a very different approach called Spectral Meta Learner (SML) (Parisi et al., 2014). SML is an unsupervised ensemble method designed for two-class classification problems. Briefly, it takes a matrix of predictions, *P*, where each row corresponds to different samples being classified and the columns correspond to different methods. Accordingly, each matrix element *P*_*ij*_ is the class (0 or 1) assigned to sample *i* by method *j* Under the assumption of conditional independence of methods given class labels, SML can estimate the balanced accuracy of each classifier in a totally unsupervised manner using only the prediction matrix *P*. The algorithm then uses this information to construct an ensemble classifier in which the contribution of each classifier is proportional to its estimated performance (balanced accuracy). The module identification problem is an unsupervised problem by its nature and we applied the SML algorithm as a new way for constructing consensus modules. For each method *m* and network *k*, we created a vector of prediction *P*_*mk*_, of size 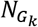 by 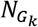, where 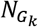 is the number genes in network as follows:

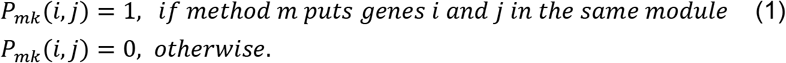 For each network, we constructed the prediction matrix *P*with each column *P*_*m*_ defined as above. We then provided this matrix as input to the SML algorithm. The SML algorithm outputs a consensus matrix, which assigns a weight between each pair of genes. We found that SML did not perform well in the context of this challenge, likely because the underlying assumption of SML is that top-performing methods converge to similar predictions, which was not the case here (see Figs. 3 and **S3**). **(E)** Pairwise similarity of networks. The upper triangle of the matrix shows the percent of shared links (the Jaccard index multiplied by 100) and the lower triangle shows the fold-enrichment of shared links compared to the expected number of shared links at random. The two protein-protein interaction networks are the two most similar networks, yet they have only 8% shared edges. Of note, a recent study has found similarly low overlap between protein interaction networks from different sources, suggesting that these molecular maps are still far from complete (Huang et al., 2018).

**Figure S3:**
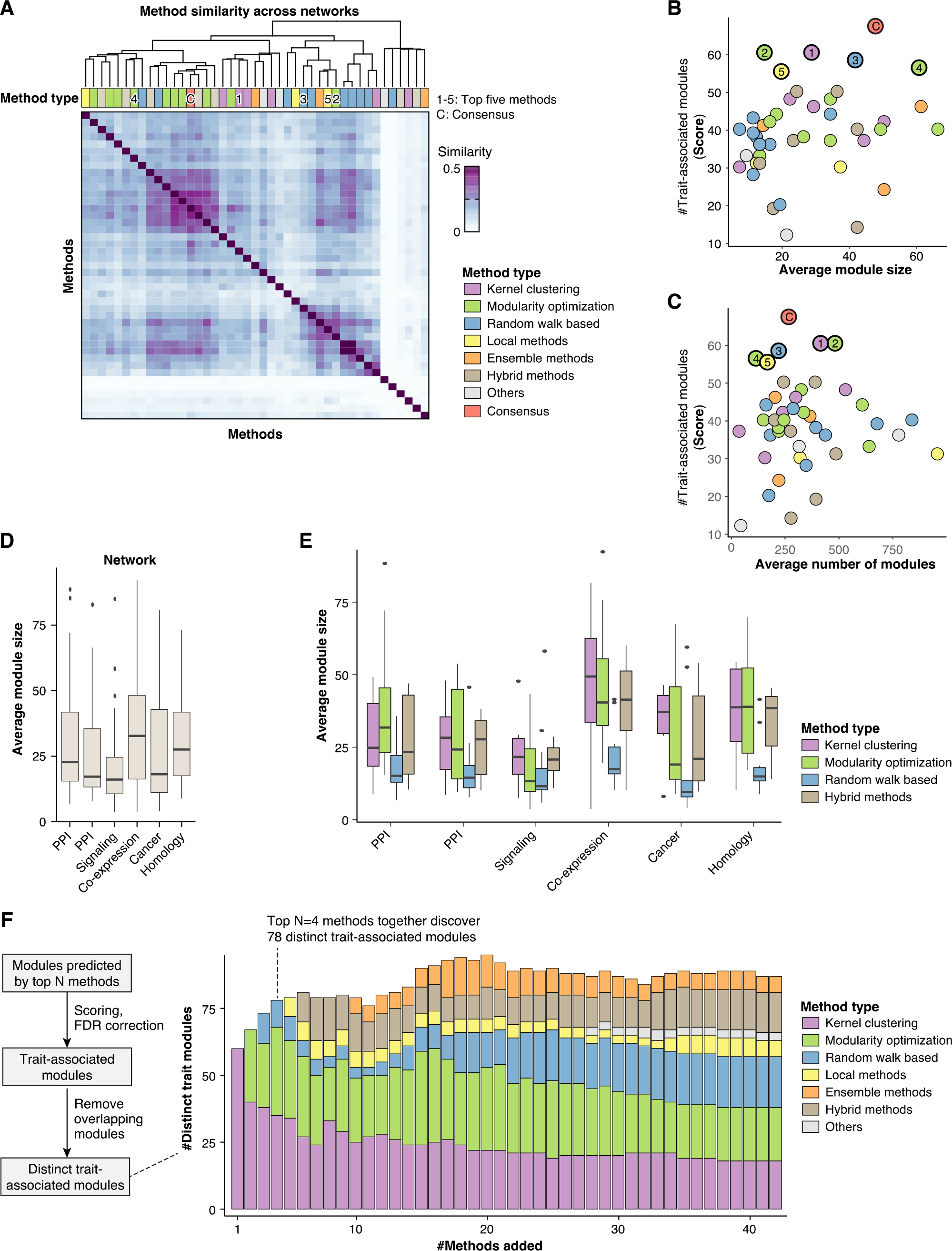
Complementarity of Module Identification Methods. All panels show results for single-network module identification methods (Sub-challenge 1). **(A)** Pairwise similarity of module predictions from different methods, averaged over all networks. Similarity was computed based on whether the same genes were clustered together by the two methods (see Methods). The resulting similarity matrix was hierarchically clustered using Ward’s method. The top row shows the method type. The top five methods (1-5) and the consensus (C) are highlighted. The top methods did not converge to similar module predictions (they are not all grouped together in the hierarchical clustering). Related to Fig. 3, which shows similarity of module predictions from individual networks. **(B)** Average module size versus score for each method. The x-axis shows the average module size of a given method across the six networks. The y-axis shows the overall score of the method. Top teams (highlighted) produced modules of varying size, i.e., they did not converge to a similar module size during the leaderboard round. There is no significant correlation between module size and score (p = 0.13), i.e., the scoring metric did not generally favor either small or large modules. **(C)** Average number of modules versus score for each method. The x-axis shows the average number of submitted modules across networks for a given method, and the y-axis shows the corresponding score. The top five teams (highlighted) submitted a variable number of modules (between 103 and 470 modules, on average, per network). There is no significant correlation between the number of submitted modules and the obtained score (p = 0.99), i.e., the scoring metric was not biased to generally favor either a small or high number of submitted modules (see Section “Scoring Metric” in the Methods for details). **(D)** Comparison of module sizes between networks. For each network, the boxplot shows the distribution of average module sizes of the 42 challenge methods. On average, modules were smallest in the signaling network and largest in the co-expression network. **(E)** Comparison of module sizes between method types. For each network, boxplots show the distribution of average module sizes for kernel clustering, modularity optimization, random-walk-based, and hybrid methods (the remaining categories are not shown because they comprise only three methods each). Note that teams tuned the resolution (average module size) of their method during the leaderboard round. The variation in module size between different method categories and networks suggests that the optimal resolution is method-and network-specific. For example, teams using random-walk-based methods tended to choose a higher resolution (smaller average module size) than teams using kernel clustering or modularity optimization methods. **(F)** Number of distinct trait-associated modules recovered by the top *K* methods. Given the top *K* methods, we considered the set including all modules predicted by these methods and scored them with the same pipeline as used for the individual methods in the challenge. We then evaluated how many “distinct” trait-associated modules were recovered by these methods. Distinct modules were defined as modules that do not show any significant overlap among each other. Overlap between pairs of modules was evaluated using the hypergeometric distribution and called significant at 5% FDR (Benjamini-Hochberg adjusted p-value < 0.05). From the set of trait-associated modules discovered by the top *K* methods, we thus derived the subset of distinct trait-associated modules (when several modules overlapped significantly, only the module with the most significant GWAS p-value was retained). Although the resulting scores (number of distinct trait-associated modules) cannot be directly compared with the challenge scores (because module predictions had to be strictly non-overlapping in the challenge), it is instructive to see how many distinct trait modules can be recovered when applying multiple methods. The stacked bars (colors) further show how many of the distinct trait modules are contributed by each method category. The number of distinct trait modules is not monotonically increasing as more methods are added because the larger sets of modules also increase the multiple testing burden of the GWAS scoring. The top four methods together discover 78 distinct trait-associated modules. Relatively little is gained by adding a higher number of methods.

**Figure S4:**
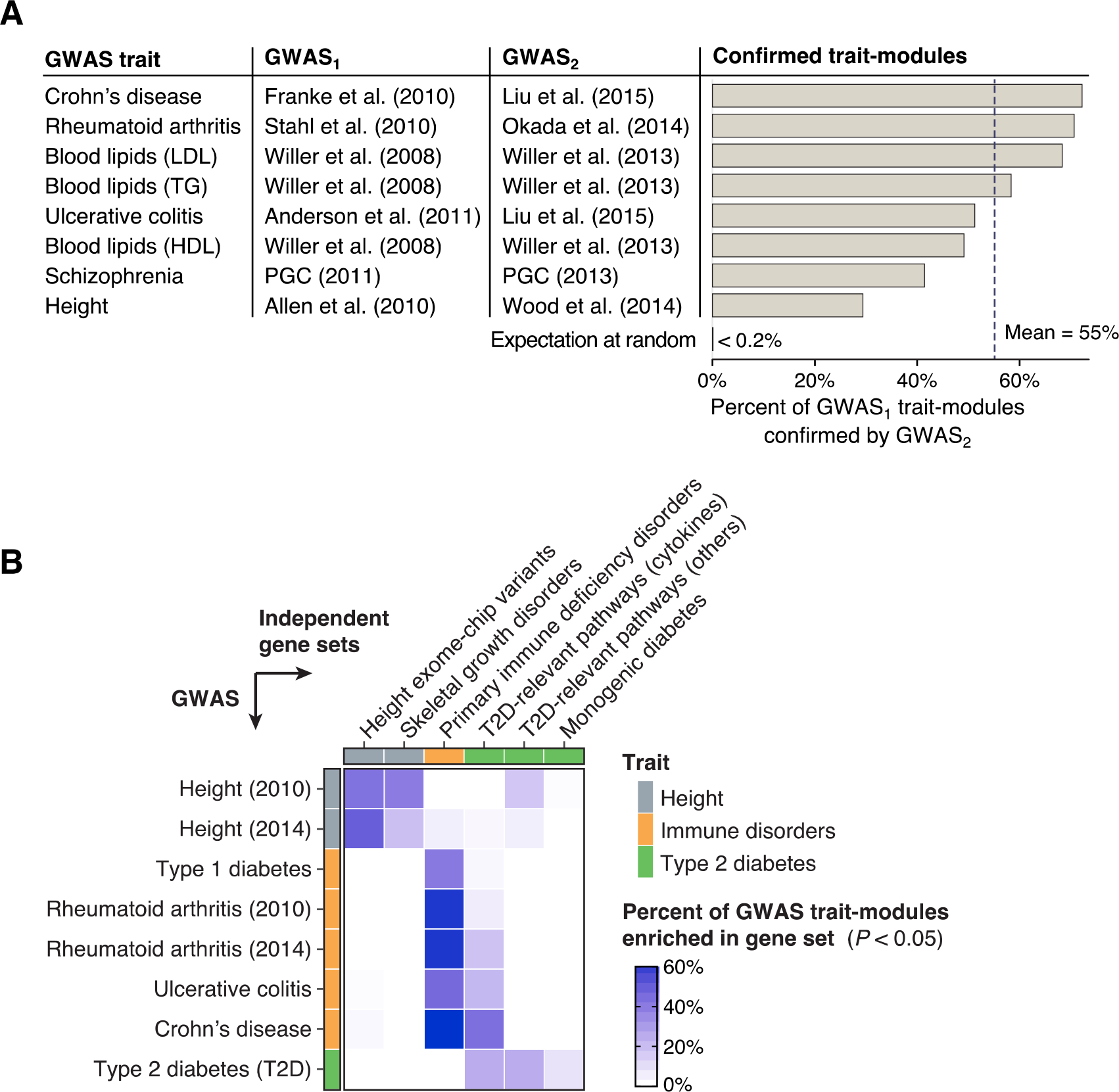
Support of Trait Modules in Diverse Datasets. **(A)** Pairs of older (lower-powered) and more recent (higher-powered) GWASs used for the evaluation of module-based gene prioritization in Fig. 5C. The first column gives the trait and the second and third columns indicate the approximate cohort sizes of the respective GWASs. The bar plot shows the percentage of trait-associated modules from the first GWAS that are also trait-associated modules in the second GWAS. At the bottom, the expected percentage of confirmed modules at random is shown (i.e., assuming the trait-associated modules in the second GWAS were randomly selected from the set of predicted modules). **(B)** Enrichment of trait-associated modules in six curated gene sets from three recent studies. The first two gene sets were taken from Marouli et al., (2017) and correspond to genes comprising height-associated ExomeChip variants and genes known to be involved in skeletal growth disorders, respectively. The third gene set was taken from de Lange et al., (2017) and corresponds to genes causing monogenic immunodeficiency disorders. Lastly, three gene sets relevant for type 2 diabetes (T2D) were taken from and correspond to genes in literature-curated pathways that are believed to be linked to T2D (we distinguished between genes in cytokine signalling pathways and other pathways) and genes causing monogenic diabetes. We then considered corresponding GWAS traits in our hold-out set, namely height, all immune-related disorders, and T2D. We then tested all modules associated with these GWAS traits for enrichment in these six external gene sets. Enrichment was tested using the hypergeometric distribution and p-values were adjusted to control FDR using the Benjamini-Hochberg method. The heatmap shows for each GWAS (row) the fraction of trait-associated modules that significantly overlap with a given gene set (column). It can be seen that modules associated with a given trait predominantly overlap the external gene sets that are expected to be relevant for that trait.

**Figure S5:**
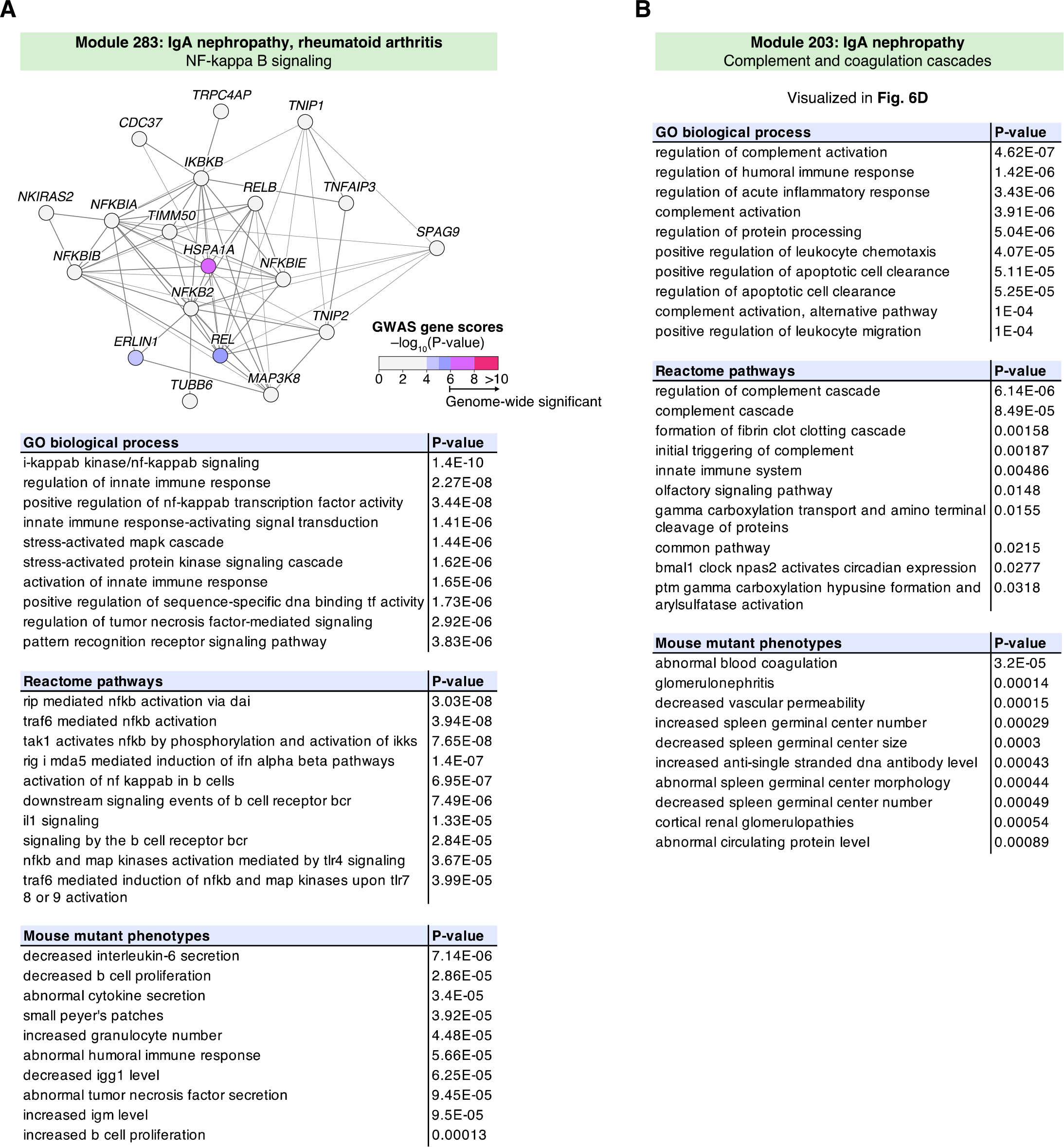
Modules Associated with IgA Nephropathy. The top ten enriched GO biological processes, Reactome pathways and mouse mutant phenotypes are shown for two IgA nephropathy (IgAN) associated modules. P-values were computed using the non-central hypergeometric distribution, see Methods. **(A)** IgAN-associated module identified using the consensus analysis in the InWeb protein-protein interaction network. The module comprises immune-related NF-κB signaling pathways. Enriched mouse mutant phenotypes for module gene homologs include perturbed immunoglobulin levels (IgM and IgG1). The module implicates in particular the NF-κB subunit *REL* as a candidate gene. The *REL* locus does not reach genome-wide significance in current GWASs for IgAN but is known to be associated with other immune disorders such as rheumatoid arthritis. **(B)** Enriched annotations for the IgAN-associated module shown in Fig. 6D. The module comprises complement and coagulation cascades. The top two enriched mouse mutant phenotypes are precisely “abnormal blood coagulation” and “glomerulonephritis”. See main text for discussion.

**Table S1:** Collection of GWAS Datasets used for the Challenge. The table lists the GWAS datasets used for the module scoring. The first column indicates whether the GWAS was used during the “leaderboard” or “final” evaluation phase. The five GWAS listed in the end (“extra”) were not used for the scoring as they were added to the collection after the challenge. The PASCAL gene scores for all GWAS are available for download from the challenge website (file names are given in the last column). The original GWAS SNP summary statistics can be downloaded individually from the indicated sources or we can share the complete collection upon request.

**Table S2:**
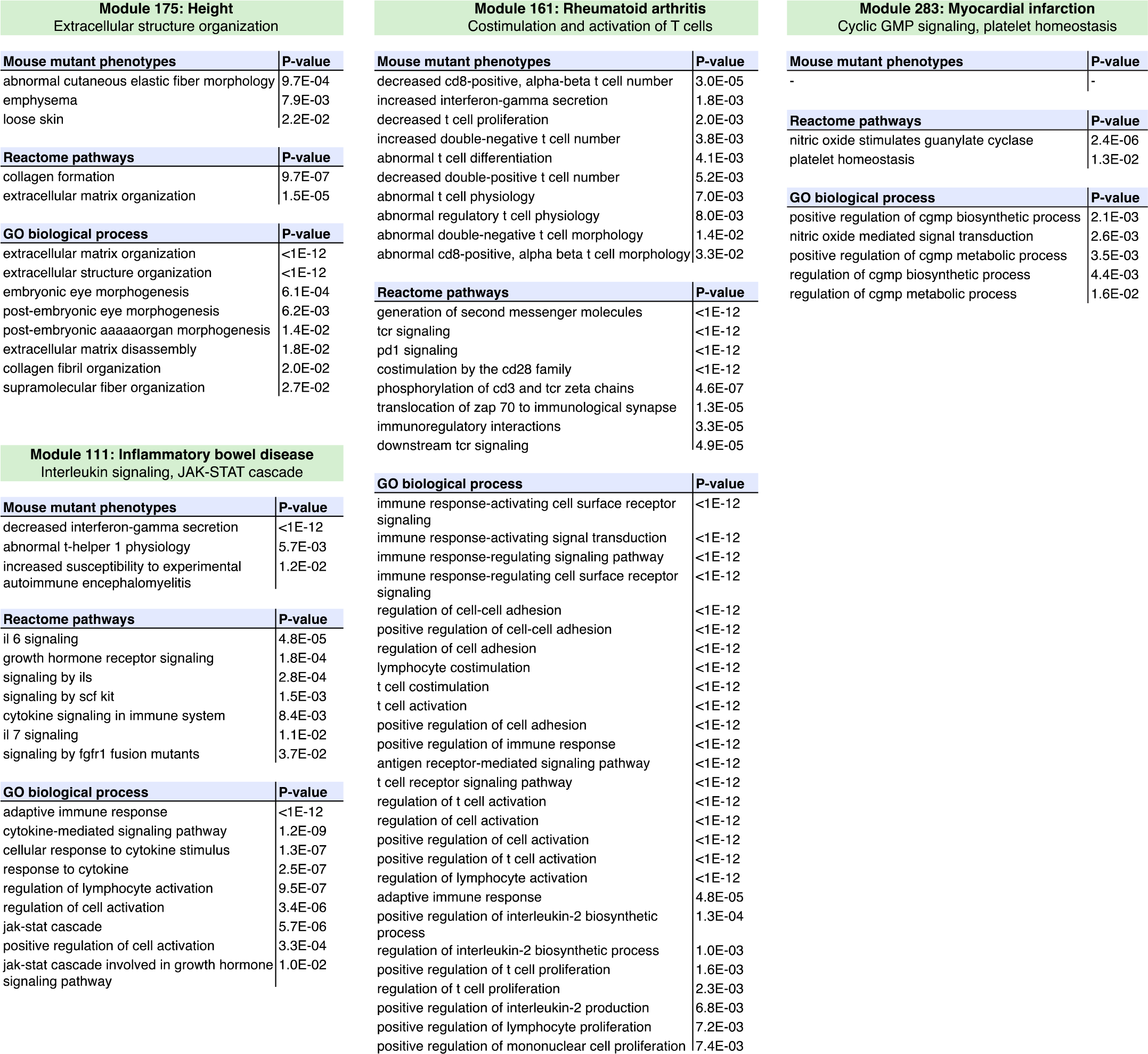
Functional Enrichment for Example Modules. Enrichment p-values for mouse mutant phenotypes, Reactome pathways and GO biological processes are shown for four example modules discussed in the main text (Figs. 5 **and** 6). P-values were computed using the non-central hypergeometric distribution and adjusted using the Bonferroni method (Methods). Results for the remaining trait-associated modules from the consensus analysis in the STRING protein interaction network are shown in **Table S4**. Functional enrichment analysis for additional pathway databases and modules from all methods and networks are available on the challenge website.

**Table S3:** Overview of Consensus Trait-modules in the STRING Network. Overview of all 21 trait-associated consensus modules in the STRING protein-protein interaction network. The first three columns give the module ID, the trait type, and the specific GWAS trait that the module is associated to. We tested all modules for enrichment in GO annotation, mouse mutant phenotypes, and other pathway databases using the non-central hypergeometric test (Methods). The putative function of each module based on this enrichment analysis is summarized in the fourth column (see Figs. 5, 6 and Tables S2, **S4** for details). Two thirds of the modules have functions that correspond to core pathways underlying the respective traits, while the remaining modules correspond either to generic pathways that play a role in diverse traits or to pathways without an established connection to the considered trait or disease. Only pathways with a well-established link to the trait were considered core pathways. Generic pathways, such as cell-cycle-related or epigenetic pathways, were not considered core pathways because they are relevant for many traits and tissues, making them more difficult to target therapeutically. For example, modules 77 and 109 are both associated with schizophrenia and comprise pathways related to epigenetic gene silencing and nucleosome organization, respectively. Although there is evidence that epigenetic mechanisms may play a role in schizophrenia, we considered this to be a generic pathway.

|  | ID | Trait type | GWAS trait | Module function / pathways | References |
| --- | --- | --- | --- | --- | --- |
| Modules | 13 | Psychiatric | Psychiatric (cross-disorder) | Neural development, chemical synaptic transmission | Table S4 |
|  | 21 | Cardiovascular | Cardiovascular disease | Lipid and steroid hormone metabolism | Table S4 |
|  | 24 | Blood lipids | Total cholesterol | Cholesterol homeostasis | Table S4 |
|  | 36 | Inflammatory | Rheumatoid arthritis | Interferon signaling, antigen processing | Table S4 |
|  | 65 | Blood lipids | LDL cholesterol | Protein ubiquitination | Table S4 |
|  | 72 | Psychiatric | Neuroticism | Mitotic cell cycle | Table S4 |
|  | 77 | Psychiatric, anthropometric | Schizophrenia, BMI | Chromatin organization, epigenetic gene silencing | Table S4 |
|  | 80 | Glycemic | Fasting glucose | Glycerophospholipid and fatty acid metabolism | Table S4 |
|  | 109 | Psychiatric, anthropometric | Schizophrenia, BMI | Nucleosome organization | Table S4 |
|  | 111 | Inflammatory | Inflammatory bowel disease | Interleukin signaling, JAK-STAT cascade | Fig. 6B, Table S2 |
|  | 126 | Anthropometric | Overweight | Activation of MAPK and JNK cascades | Table S4 |
|  | 161 | Inflammatory | Rheumatoid arthritis | Costimulation and activation of T cells | Fig. 6A, Table S2 |
|  | 175 | Anthropometric | Height | Extracellular structure organization | Fig. 5, Table S2 |
|  | 248 | Anthropometric | Overweight | Lipid and insulin signaling, B cell activation | Table S4 |
|  | 262 | Anthropometric | Height | Pre-NOTCH transcription and translation | Table S4 |
|  | 270 | Inflammatory | Ulcerative colitis | MHC class II antigen presentation | Table S4 |
|  | 271 | Inflammatory | Inflammatory bowel disease | Interleukin signaling, regulation of JAK-STAT cascade | Table S4 |
|  | 283 | Cardiovascular | Myocardial infarction | Cyclic GMP signaling, platelet homeostasis | Fig. 6C, Table S2 |
|  | 294 | Anthropometric | Body mass index (women) | Phospholipid metabolism and transport | Table S4 |
|  | 323 | Inflammatory | Crohn's disease | TNF-mediated signaling | Table S4 |
|  | 326 | Blood lipids | LDL cholesterol | Plasma lipoprotein particle clearance | Table S4 |

**Table S4:** Functional Enrichment of Consensus Trait Modules. For each of the 21 consensus trait-modules shown in Table S3, all categories with a Bonferroni-corrected P-value below 0.05 are listed (Methods). Only results for mouse mutant phenotypes, Reactome pathways and GO biological process annotations are included for brevity. Full results including all tested pathway databases and all challenge modules are available on the challenge website.

